# Conservative Significance Testing of Tripartite Interactions in Multivariate Neural Data

**DOI:** 10.1101/2022.02.07.479415

**Authors:** Aleksejs Fomins, Yaroslav Sych, Fritjof Helmchen

## Abstract

An important goal in systems neuroscience is to understand the structure of neuronal interactions, frequently approached by studying functional relations between recorded neuronal signals. Commonly used pairwise metrics (e.g. correlation coefficient) offer limited insight, neither addressing the specificity of estimated neuronal interactions nor potential synergistic coupling between neuronal signals. Tripartite metrics, such as partial correlation, variance partitioning, and partial information decomposition, address these questions by disentangling functional relations into interpretable information atoms (unique, redundant and synergistic). Here, we apply these tripartite metrics to simulated neuronal recordings to investigate their sensitivity to impurities (like noise or other unexplained variance) in the data. We find that all considered metrics are accurate and specific for pure signals but experience significant bias for impure signals. We show that permutation-testing of such metrics results in high false positive rates even for small impurities and large data sizes. We present a conservative null hypothesis for significance testing of tripartite metrics, which significantly decreases false positive rate at a tolerable expense of increasing false negative rate. We hope our study raises awareness about the potential pitfalls of significance testing and of interpretation of functional relations, offering both conceptual and practical advice.

**Author Summary:** Tripartite functional relation metrics enable the study of interesting effects in neural recordings, such as redundancy, functional connection specificity and synergistic coupling. However, common estimators of such relations are designed for pure (e.g. non-noisy) signals rare for such recordings. We study the performance of tripartite estimators using simulated impure neural signals. We demonstrate that permutation-testing is not a robust procedure for inferring ground truth interactions from studied estimators. We develop an adjusted conservative testing procedure, reducing false positive rate of studied estimators for impure data. Besides addressing significance testing, our results should aid in accurate interpretation of tripartite functional relations and functional connectivity.

## 1 Introduction

Recent advances in brain recording techniques enable simultaneous acquisition of multiple neuronal signals. Examples are single-cell population recording techniques, such as multielectrode arrays [72] or two-photon calcium imaging [18], as well as multi-regional population-average recording techniques, such as wide-field imaging [36], multi-fiber photometry [73], EEG [57], MEG [19] or fMRI [44]. An important stepping stone to understand neural coding is the ability to robustly infer and interpret possible functional relations between multivariate signal components, be it single neurons or population-averaged regional signals. At first glance, the procedure may appear as simple as computing a standard relational metric, such as Pearson’s correlation coefficient, followed by reporting the pairs of signals with high or low coefficient values. However, a finer inspection reveals several pitfalls of such an approach. The aim of this paper is to illuminate one such pitfall, discuss its implications and propose a solution.

Specifically, we address the negative effects of data impurities on the robustness of functional relation estimates.

Functional relations can be defined via a model-based approach. A general model will attempt to explain one of the signals, known as the dependent variable (or simply the target), by means of other signals, known as the independent variables (or sources, or predictors). The special case of considering a single source is covered by the the well-studied fields of pairwise functional connectivity [33] and effective connectivity [39]. Introduction of multiple sources enables the study of interesting higher-order effects, such as confounding effects on pairwise connections as well as synergistic effects between sources. Here, we focus our attention on two source variables, i.e. on tripartite metrics. The use of tripartite functional relations in addition to functional connectivity may pave the way towards causal interpretations of neuronal recordings [64], albeit not without shortcomings [55] or additional research. While considering a larger number of source variables may be considered in principle [80], it is less feasible. The number of possible types of higher-order relations grows exponentially with the number of variables, as does the data size required for robust estimation of such relations.

A pair of source variables *X* and *Y* may contain information about a target variable *Z* in 4 distinct ways [80], called information atoms (see fig. 1). We aim to reveal how well different metrics framed in this formalism can recover ground truth information in simulated multivariate recordings. Two concepts that make such estimation challenging are redundancy and impurity. We will now introduce them.

**Figure 1:**
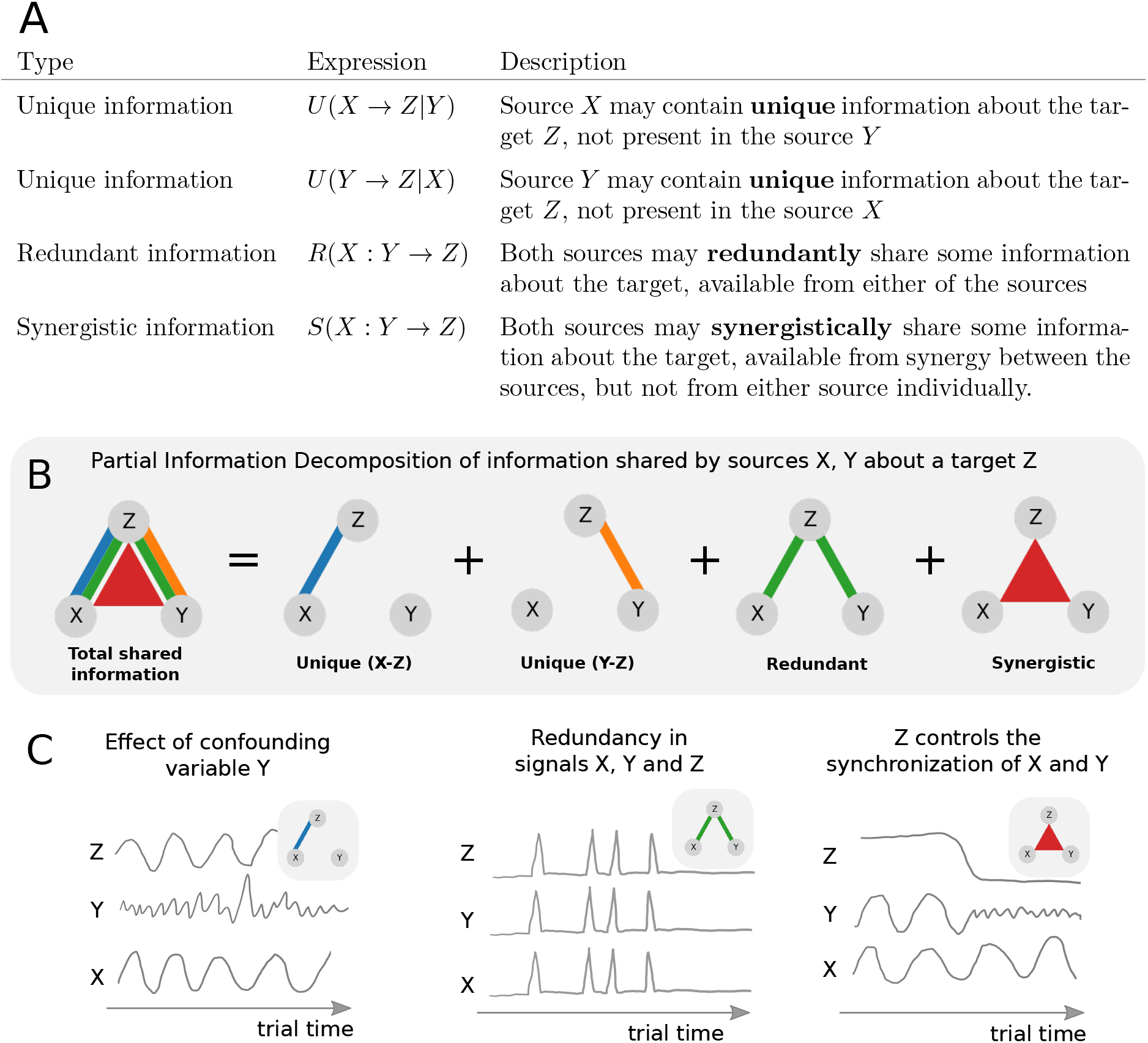
A) Four information atoms of Partial Information Decomposition (PID) [80]. *X*, *Y* and *Z* are three recorded variables (e.g. neuronal signals). Here, *X* and *Y* are the independent (source) variables, and *Z* is the dependent (target) variable. B) Sketch of PID. Sketches of this form will be employed throughout this paper. The colors will always denote the corresponding information atoms *U* (*X* → *Z*|*Y*), *U* (*Y* → *Z*|*X*), *R*(*X* : *Y* → *Z*), *S*(*X* : *Y* → *Z*). The width of individual lines or triangles qualitatively indicates the magnitude of the effect. In this plot, all information atoms are shown with maximal magnitude for reference. C) Example questions about tripartite relations that may be of interest in neuroscience. Left: Is the functional connection between *X* and *Z* specific with respect to the confounding variable *Y* ? Middle: Are *X*, *Y* and *Z* redundantly encoding the same information? Right: Could *Z* control synchronization between *X* and *Y* , (for example, if *X* and *Y* control forelimbs and hindlimbs respectively, and *Z* determines if the animal is currently running or resting). Note: the three sketches are made as function of time for illustrative purposes only. In principle, information atoms can be computed across any data dimension. Here, we compute information atoms across trials.

We first consider redundancy. A common method for studying linear relations between source and target variables is Multi-way ANalysis Of VAriance (ANOVA) [37]. It provides information about the overall goodness of fit of a model, as well as on the expected magnitude and significance of individual coefficients. While ANOVA is known to provide robust estimates of coefficient significance when the source variables are mostly unrelated [4], it fails to do so when the source variables are related. This phenomenon is known as *multicollinearity* [27] in statistics literature and as *redundancy* in neuroscience [46]. In case of redundancy, a broad range of parameter value combinations may result in an optimal model fit. Hence, multiple different parameter combinations may be indistinguishable to the fitting procedure. In such case, ANOVA will arbitrarily report some parameter values resulting in a good fit, with unreliable estimates of parameter significance [27]. This effect is undesirable, as we ultimately want to know the importance and specificity of individual sources as predictors. Importantly, high redundancy is common in both single-neuron recordings [35] and in multi-regional population-average recordings [36, 74], and thus needs to be accounted for.

Next we consider impurity. Neuronal recordings frequently do not directly access the neuronal variables of interest. Apart from instrumental noise, observables may be corrupted by various other factors including imperfect knowledge of the properties of the signal proxy (e.g. calcium indicator or BOLD fMRI response), contamination by neuropil fluorescence signals or cross-talk, and heart-beat or movement induced artifacts. Although such impurities are typically acknowledged in the experimental literature, they often are overlooked in statistical analyses such as functional connectivity estimation. Consider a simple linear model

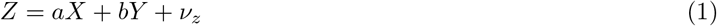

where *Z* is the target variable, *X* and *Y* are the source variables, *a* and *b* are the corresponding coefficients and *ν*_*z*_ is the residual error. In this case, *Z* is corrupted by the additive error *ν*_*z*_. While it is tempting to call *ν*_*z*_ noise, it is better described as impurity, and its variance as unexplained or residual variance. While part of it may be due to experimental limitations as described above, impurity may also arise due to other sources that have not been observed in the experiment. For example, the mood of a cat may be affected by weather and the quality of their meal, but also by the amount of petting they have received. An optimal model that includes all of these sources will have lower residual variance in explaining the cat’s mood than an optimal model that does not include petting. Calling the residual variance in the latter model noise would be misleading as it correlates with petting, which could have been recorded and taken into account. Such scenarios are common in neuroscience. For example, a population-average signal may represent multiple distinct neuronal sub-populations with different functional connectivity, such that only part of the observed signal correlates with the signal of interest (e.g. the activity in another brain area). Similarly, an individual neuron may integrate multiple inputs, of which not all are recorded. Impurity of observables in terms of residual variance thus does not solely reflect limitations of the measurement techniques, but also the incompleteness of observing all relevant sources.

Direct, i.e. pure, access to source variables of interest is also not a given. For example, the source variables may contain additive impurities *ν*_*x*_ and *ν*_*y*_ of similar origins as described above for the target variable. In general, all three variables may contain impurities (fig. 2). For simplicity, we will only consider additive errors, although in general the relation may be more complex. We will denote the underlying neuronal variables with an asterisk (e.g. *X*\*) and the corresponding observables without one (e.g. *X*). The impurities *ν*_*x*_, *ν*_*y*_ and *ν*_*z*_ are assumed to be statistically independent in this work.

**Figure 2:**
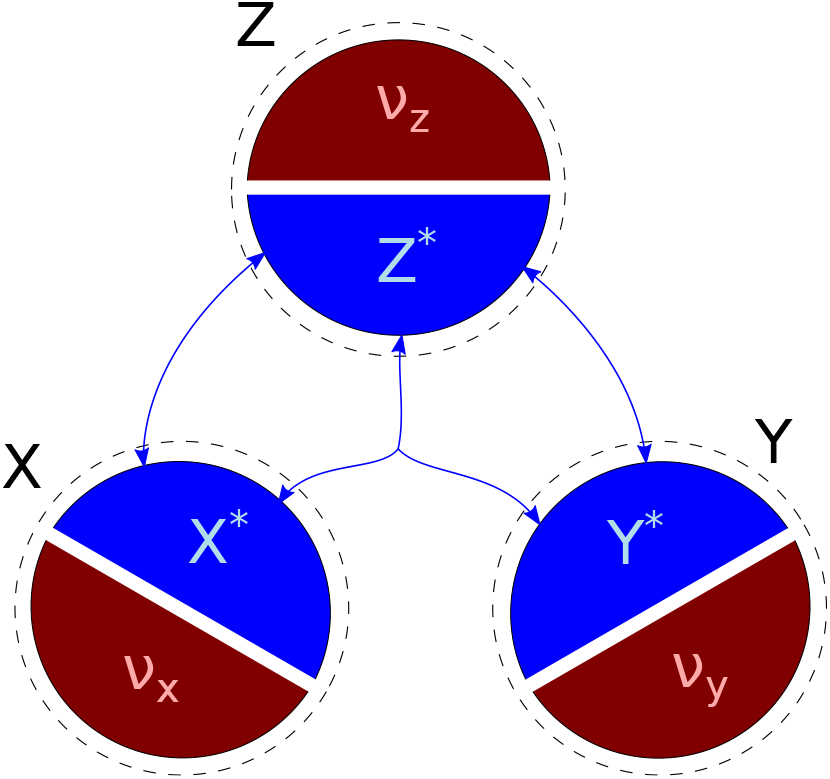
Impurity in neuronal observables. A typical aim is the estimation of information atoms (blue arrows) between neuronal signals of interest (blue areas *X*\*, *Y* * and *Z*\*) underlying the recorded data. However, the observables the experimenter has access to (black areas *X*, *Y* and *Z*) typically are not the pure signals of interest. In the simplest case considered here, observables are corrupted by additive impurities (red *ν*_*x*_, *ν*_*y*_ and *ν*_*z*_). Blue arrows in the middle indicate tripartite interaction effects between the signals of interest (i.e. synergy).

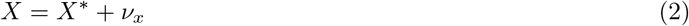

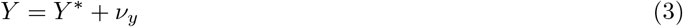

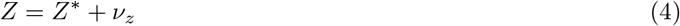

To describe the purity level of the relation between the observable and the underlying variable of interest we define the *impurity fraction* (see section 2.2.2). Impurity fraction is a number between 0 and 1, where 0 denotes a pure signal with no residual errors, and 1 denotes a signal consisting only of residual errors. It is similar to signal-to-noise ratio (SNR) that is commonly used in signal theory. However, SNR does not cover the case of 100% signal, which we find interesting to consider. Further, we believe that the word noise in SNR may be misleading for the reasons outlined above.

Many metrics are designed to estimate functional relations (functional connectivity or information atoms) between pure variables. The presence of impurities, especially in source variables, frequently results in violation of the assumptions of these metrics, and thus may produce spurious findings. In statistics and econometrics, models aware of potential source variable impurities are known as errors-in-variables models [38]. For example, the term regression dilution [43] describes the effect that basic linear regression will increasingly underestimate the absolute value of the regression coefficient with increasing impurity in the source variables. In the neuroscience community, we believe that detrimental effects of impurities on multivariate estimators are less well known, motivating us to attract attention to these effects here.

Having introduced redundancy and impurity, we will now outline the scope of this study. Our specific aims are to present metrics designed to disentangle individual functional relations between triplets of variables in the presence of redundancy, to computationally test whether these metrics are robust to impurities in source and target variables, and to propose and discuss potential improvements. We focus on three existing metrics: Partial Correlation (PCorr) [30], Variance Partitioning (VP) [12], and Partial Information Decomposition (PID) [80]. Precise definitions of these metrics are given in the methods section. Partial Correlation has been used in neuroscience to study the specificity of functional connections between neurons [25] and fMRI voxels [53, 31]. A recent study [41] proposed a test for PCorr taking signal autocorrelation into account, which is of high relevance for neuronal signal proxies such as calcium indicator or fMRI BOLD signals. Variance Partitioning, previously introduced in ecological analysis [12, 59, 11], was recently used to study unique and redundant feature encoding in human fMRI recordings [48, 45]. The original method is based on decomposing the variance explained by a combination of sources, obtaining unique and redundant explained variances. In this paper, we extend this methodology by also including quadratic synergistic terms, thus making VP comparable to PID described below. VP is strongly related to Partial R-Squared (also known as Partial F-Test), which is a popular metric because it allows for quantitative comparison of two linear models explaining the same target variable. In neuroscience, among other fields, it has been used to compare models of hemodynamic response in fMRI [2], shape-selectivity in V4/IT [13], reaction time in working-memory tasks [28], fatigue in multiple sclerosis [56], and neuronal correlates of minimal conscious state [62]. Partial Information Decomposition is the most recent of the metrics. While it has been actively developed by the information-theoretic community since a decade [80], it has been rapidly gaining popularity in neuroscience in the last few years. For example, PID has been used to demonstrate a relationship between synergy and feed-back information flow in mouse organotypic cultures [69], to show significant synergy between somatic and apical dendritic output of L5b pyramidal neurons and its relationship to activation of dendritic *GABA*_*B*_ receptors in rat S1 slices [68], and to estimate unique contributions of acoustic features of speech to BOLD responses in humans [22, 23]. Further, it has been used to explore the structure of simulated input-driven recurrent network models [16] and artificial generative neuronal networks [75]. We believe that PID will be increasingly applied in coming years, especially in studies addressing non-linear confounding effects, the specificity of functional relations, and synergistic encoding.

In the following, we ask whether these metrics are sensitive and specific in detecting individual interactions in simulated data with known ground truth. All of the tested metrics turn out both significant and specific for model data with pure (error-free) source variables. However, addition of even small impurities to the source variables damages the specificity of the metrics when permutation-tested. As a partial remedy, we propose a null hypothesis which corrects the bias introduced by the impurity. Compared to permutation-testing, this approach significantly reduces the false positive rate at the expense of increasing the false negative rate. This approach should be beneficial in exploratory neuroscience research, aiming to preserve robustness of the stronger findings at the expense of losing some of the weaker ones.

## 2 Methods

Let us consider the following scenario (fig. 3A): A test subject (e.g. a mouse or a human) performs a temporally structured behavioural task while brain activity is simultaneously recorded via three neuronal observables *X*, *Y* and *Z*. Depending on the recording method, the observables may represent single-cell activity or regional bulk activity, pooled across multiple neurons. The test subject repeats the task over a set of trials, which are of equal duration and assumed to be independent and identically distributed (i.i.d.). In total, *N* = *N*_*trial*_*N*_*time*_ datapoints are recorded for each observable, where *N*_*trial*_ is the number of trials and *N*_*time*_ is the number of time steps in a single trial. We want to understand how the signals *X* and *Y* and their interactions may be related to *Z*. More precisely, the aim is to quantify the functional relations between two source signals, *X* and *Y* , and the target signal *Z* (by means of information atoms) and to evaluate how they change over trial time. Here, we study information atoms across trials for a fixed time point. This approach satisfies the i.i.d. requirement of information atom estimators used in the paper. The process can be repeated for every time step individually, which allows to build up the temporal dynamics of the information atoms. Thus, the problem of studying time-dependent evolution of functional relations between three neuronal observables can be reduced to the problem of estimating the information atoms from *N*_*trial*_ i.i.d simultaneous samples of the random variables *X*, *Y* and *Z*. Possible extensions of the above assumptions are addressed in the discussion.

**Figure 3:**
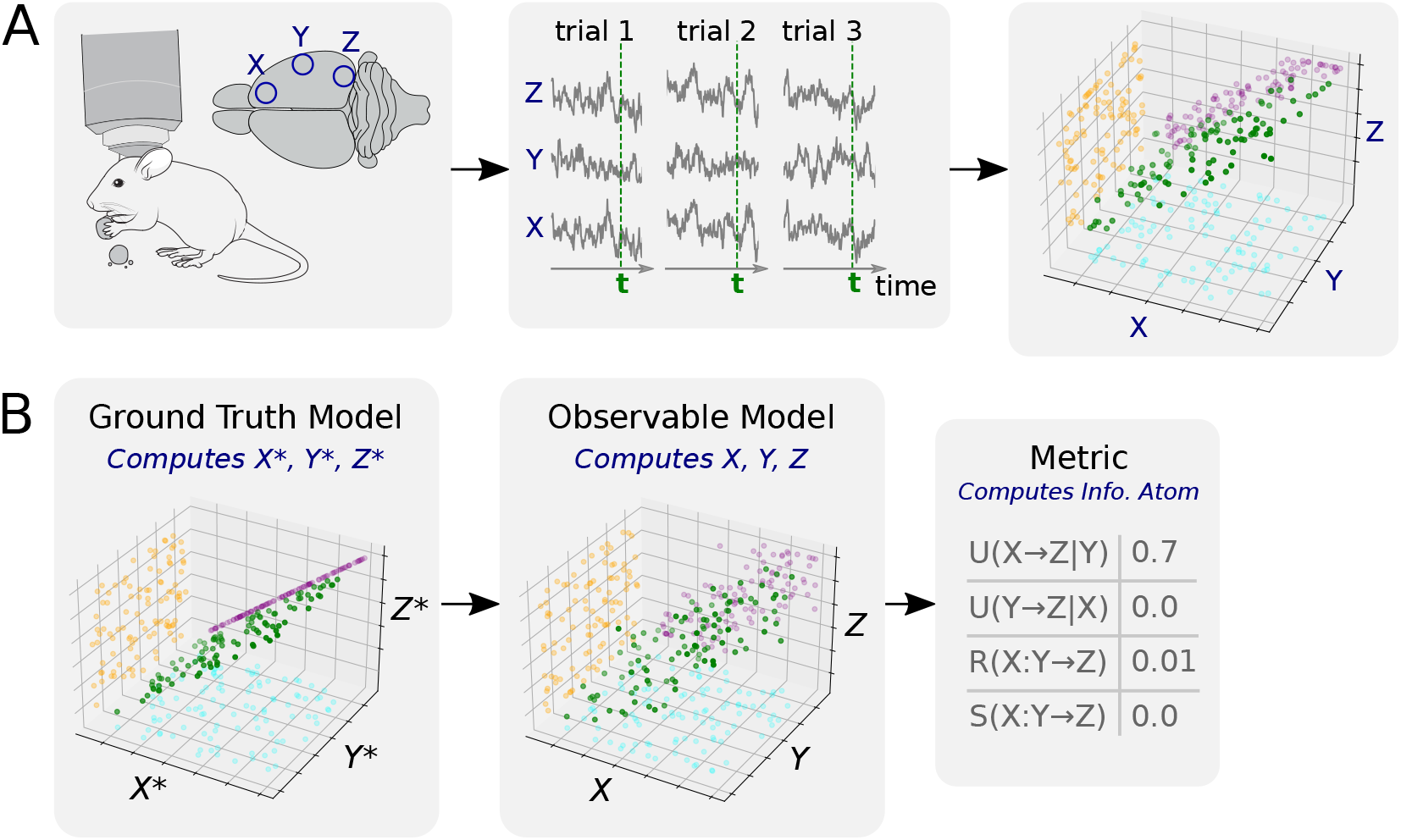
A) A thought experiment setup. Left: Multivariate neuronal signals are recorded in a behaving test subject (courtesy to SciDraw). Middle: Neuronal signals *X*, *Y* and *Z* are observed during *N*_*tr*_ trials with the same duration *T* and are plotted as a function of trial time for three example trials. Green vertical lines indicate a sample time step at which the analysis is performed. Right: 3D scatter plot of *X*, *Y* and *Z* across trials sampled at the fixed time step *t* (green). 2D projections indicate that *X* correlates to *Z* (purple), while *Y* is uncorrelated to either *X* or *Z*. B) A sketch of the simulation procedure. First the ground truth model is used to generate multiple samples of the ground truth variables *X*\*, *Y* *, *Z*\*. Then, the observable model adds impurity to the data, producing observables *X*, *Y* , *Z*. Finally, the metric is used to compute information atoms for the given data sample.

In the following, we first present the three metrics that we used to estimate the tripartite functional relations. Second, we introduce three ground truth models that we used to simulate the ground truth variables at a fixed time step over trials. Third, we present observable models that we used to obtain the observable variables from the ground truth variables by adding impurities. Finally, we explain the testing procedure we used test the significance of individual information atoms. The summary of the simulation procedure is given in fig. 3B.

### 2.1 Metrics for Tripartite Analysis

#### 2.1.1 Partial Correlation (PCorr)

PCorr is the Pearson’s Correlation Coefficient between two random variables *X* and *Z*, controlling for the confounding variable *Y* . The control is performed by fitting *Y* to each of *X* and *Z* using linear least squares, subtracting the fits to obtain residuals,

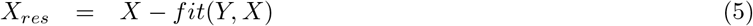

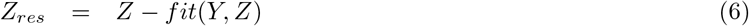

followed by computation of the Pearson’s correlation coefficient between the residuals

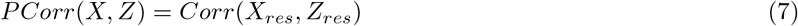

Similarly, the partial correlation *P Corr*(*Y, Z*) between *Y* and *Z* can be computed by finding and correlating the residuals of both variables with respect to *X*.

#### 2.1.2 Variance Partitioning (VP)

Partial R-Squared (*PR*^2^) is a metric generally used for quantifying the difference in performance of two linear regression models in explaining the same dependent variable. In practice, it is commonly used to evaluate the usefulness of individual independent variables. Using the three variable example, we might want to estimate the usefulness of the source *X* as predictor of the target *Z*, given another source variable *Y* . To do so, we would construct a model *f* of two variables *Z*_*f*_ = *f* (*X, Y*) and another simpler model *g* without *X*, i.e. *Z*_*g*_ = *g*(*Y*). After fitting both models, we would compute the residual sum of squares (SSR) for each model. SSR is the ”unexplained” sum of squares, calculated after the model has been fitted to and subtracted from the target.

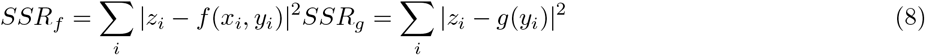

*PR*^2^ is defined as the difference of these two residual terms. Here, backslash denotes set exclusion (i.e., */X* denotes a model where *X* is excluded from the set of predictors; in this case only *Y* remains).

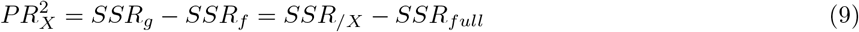

*PR*^2^ can be used to define Variance Partitioning (VP). First, a full model *F* with all of the predictors of interest is fitted to the target variable *Z*. The total sum of squares (*SST*_*Z*_) of the target variable can then be partitioned into the sum of squares explained by the model (*SSE*_*F*_) and the sum of squares of the residuals *SSR*_*F*_.

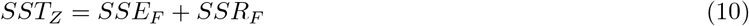

*SSE*_*F*_ can further be partitioned into non-negative parts (unique *U* , redundant *R*, and synergistic *S*) similar to those defined in Partial Information Decomposition (PID, see below). For consistency with PID, we refer to the parts of this decomposition as information atoms. We are aware that standard error does not directly measure information, and that this metric is only conceptually similar to PID.

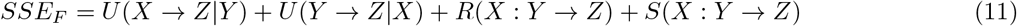

Here, VP is based on the application of *PR*^2^ to a simple quadratic interacting model with two independent variables.

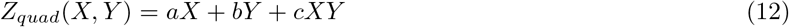

where the last term is the coupling term between *X* and *Y* , modeling their synergistic effect on *Z* . Throughout this section, we assume that means have been subtracted from both source and target variables prior to fitting. In principle, this can also be done by additionally modeling a constant term, which we drop here for simplicity. Note that the term *XY* with the coefficient *c* is also a predictor distinct from *X* and *Y* . Even though it depends on *X* and *Y* in general, it can be shown to be linearly-independent from *X* and *Y* , effectively resulting in a new predictor.

The original definition of VP [12] includes only the first two terms, i.e., modeling unique and redundant information atoms. While we are not aware of other publications using a quadratic term in this exact setting, it is commonplace to use quadratic terms to model coupling between sources in similar settings (see for example [71]). We can define unique and synergistic information atoms by the corresponding *PR*^2^, namely by the explained variance lost when excluding each of the terms in the model individually:

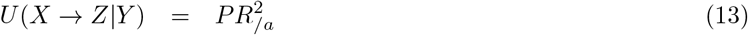

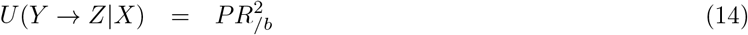

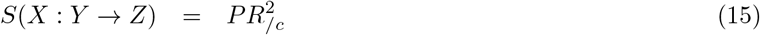

For completeness, we augment the above model by also defining the redundant information atom.

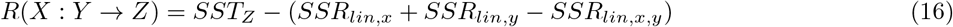

Here, *SSR*_*lin,x*_, *SSR*_*lin,y*_, and *SSR*_*lin,x,y*_ are the residual sums of squares corresponding to linear models containing only the source *X*, only source *Y* , and both sources *X* and *Y* respectively. The derivation of the *R*(*X* : *Y* → *Z*) is more technical and is thus treated in the supplementary information. In all plots, VP information atoms are normalized by *SST*_*Z*_ to obtain a dimensionless number between [0, 1]. Loosely, this number can be interpreted as the fraction of total variance explained by each information atom, although some authors have argued that this interpretation may be misleading [1]. Normalization does not affect significance testing and is done for aesthetic purposes only. Thus, we only make statements about relative values of VP information atoms, and make no statements about the interpretation of the absolute values.

Besides studying unique information atoms similar to PCorr, VP can also estimate redundant and synergistic information atoms, similar to PID discussed below. However, VP is only an approximation for relations beyond linear, and the synergistic term is only sensitive to interactions that have a non-negligible quadratic component.

#### 2.1.3 Partial Information Decomposition (PID)

PID is a non-negative decomposition of the Shannon mutual information shared by a pair of source variables *X*, *Y* and a target variable *Z* (given by the mutual information *I*(*X, Y* : *Z*)) into independent information atoms [80].

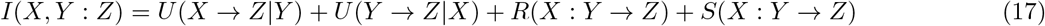

Similar to the other metrics described, unique information atoms (*U* (*X* → *Z*|*Y*) or *U* (*Y* → *Z*|*X*)) measure the information shared by the target and one of the source variables but not the other one, redundant information atoms *R*(*X* : *Y* → *Z*) measure the information shared by the target and either one of the source variables, and synergistic information atoms *S*(*X* : *Y* → *Z*) measure the information shared by the target and both the source variables but not shared by either of them independently. All information atoms are non-negative, measured in bits, and have an upper bound given by the entropy of the target *H*(*Z*). As for the other metrics, the total shared information *I*(*X, Y* : *Z*) may be significantly less than its maximum *H*(*Z*) because the sources need not be able to perfectly explain the target.

Currently available implementations of PID only work with discrete random variables, although extensions to continuous variables are actively explored [60, 66]. Thus, eq. (17) is the decomposition of the discrete (Shannon) entropy and the discrete mutual information. As PID operates on the phase-space of discrete random variables, it is in principle sensitive to arbitrary non-linear interactions between the discrete variables. However, in many cases (e.g. neural data) continuous variables are recorded. In such cases, their empirical probability distributions have to be binned so that PID is applicable to them. In this work, we focused on the analysis of continuous observables. To this end, we use equi-quantile binning, meaning that the same number of datapoints land in each bin for each variable.

Since our models contrast positive and negative values of continuous variables with symmetric distributions, we used a reasonable guess that binning these variables to 2-bin discrete distributions would work well (see models section below). It may be possible to reach similar conclusions with real data by exploring the single variable and pairwise scatter plots of the data, followed by making an ansatz about the possible analytic shape of the multi-variate probability distribution. However, in general such guesses are subjective and not robust. To study how the choice of bin number may affect information atoms and the derived results, we repeated the main analyses of this paper for different bin numbers (Supplementary Figure 18). As expected, we found that the effects of bin number are not negligible, strongly affecting false positives, especially in the redundant model. The analysis justifies our choice of 2 bins as the best possible strategy for the available data and metric. We conclude that studies interested in applying PID to continuous data should clearly report the selected bin number and also demonstrate how their results depend on small variations in bin number.

Finally, several different implementations of PID exist [9, 63, 50]. While all of the implementations agree on information-theoretic equations constraining the information atoms, they generally disagree on the definition of the redundant information atom, on the operational interpretation, as well as on whether PID should be symmetric in sources and target [63], among others (see [40] for an excellent review on this topic). Here, we chose the Bertschinger interpretation [9] and used the BROJA estimator [51, 52] from the open-source Python library IDTxl [81] to calculate PID information atoms. The main reason for this choice is the accessibility of the implementation and the fact that in Bertschinger interpretation all the information atoms remain non-negative, slightly simplifying the interpretation in our opinion. While comparison of PID implementations for multivariate analysis of neuronal data is certainly worth further studying, it is beyond the scope of this work.

### 2.2 Models

#### 2.2.1 Ground Truth Models

Here we present two linear models and one quadratic model simulating the target variable *Z*\* as a function of two source variables *X*\* and *Y* *. For non-symmetric metrics, *X*\* denotes the primary predictor of *Z*\* and *Y* * denotes the confounding predictor. Each model describes the ground truth variables *X*\*, *Y* * and *Z*\* in terms of the latent variables *T* , *T*_*x*_ and *T*_*y*_ (table 1). Each model is designed to exhibit only one of the information atoms (redundant information model **ModelR**, unique information model **ModelR** and synergistic information model **ModelS**). The purpose of this choice is to estimate false positive rates in extreme cases. We have designed a continuous-variable and a discrete-variable version of each model. Only the continuous-variable models are used in the main text of this paper. The purpose of discrete-variable models is to validate the results for the PID metric (see Supplementary Figure 17). As mentioned above, PID metrics currently are designed to work with discrete data, requiring binning of continuous variables prior to analysis. We therefore require the discrete models to estimate the extent to which the effects observed using PID metrics could be confounded by the binning procedure. In the continuous case, latent variables are modeled by standard normal variables, in the discrete case by standard Bernoulli random variables (balanced coin flips).

**Table 1:** Three ground truth models. Ground truth variables *X*\*, *Y* * and *Z*\* depend linearly on the latent variables *T* , *T*_*x*_ and *T*_*y*_. Each model has a continuous-variable and a discrete-variable version. XOR denotes the exclusive-or logical function. Information atom values of 0 and 1 are given for illustrative purposes, denoting the minimal and maximal values of the corresponding metric. 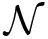 denotes a Gaussian random variable, *Ber* denotes a Bernoulli random variable.

| Shorthand | **ModelR** | **ModelU** | **ModelS** | Latent Variables |
| --- | --- | --- | --- | --- |
| Continuous Equations | $X^* = T$ | $X^* = T$ | $X^* = T_x$ | $T \sim \mathcal{N}(0, 1)$ |
| | $Y^* = T$ | $Y^* = T_y$ | $Y^* = T_y$ | $T_x \sim \mathcal{N}(0, 1)$ |
| | $Z^* = T$ | $Z^* = T$ | $Z^* = T_x T_y$ | $T_y \sim \mathcal{N}(0, 1)$ |
| Discrete Equations | $X^* = T$ | $X^* = T$ | $X^* = T_x$ | $T \sim \text{Ber}(0.5)$ |
| | $Y^* = T$ | $Y^* = T_y$ | $Y^* = T_y$ | $T_x \sim \text{Ber}(0.5)$ |
| | $Z^* = T$ | $Z^* = T$ | $Z^* = \text{XOR}(T_x, T_y)$ | $T_y \sim \text{Ber}(0.5)$ |
| $U(X \rightarrow Z\vert Y)$ | 0 | 1 | 0 | |
| $U(Y \rightarrow Z\vert X)$ | 0 | 0 | 0 | |
| $R(X : Y \rightarrow Z)$ | 1 | 0 | 0 | |
| $S(X : Y \rightarrow Z)$ | 0 | 0 | 1 | |

#### 2.2.2 Observable Models

The observable variables *X*, *Y* and *Z* represent the variables actually observed by an experimenter. They are modeled as ground truth variables with added error terms (table 2). In the continuous-variable case, the error terms are modeled as standard normal variables. The parameters *p*_*x*_, *p*_*y*_ and *p*_*z*_ are the Impurity Fractions, which are used to control the fraction of unexplained signal in the observable variables in eqs. (2) to (4) Impurity fractions are real variables in the range [0, 1]. They linearly interpolate between a pure signal perfectly explained by the ground truth model (*p* = 0), and a 100% impure signal completely unrelated to the ground truth model (*p* = 1). Changing an impurity fraction keeps the variance of the observable constant.

**Table 2:** Continuous and discrete observable models.

| Model Type | Observables | Impurity Fraction | Unexplained Variance |
| --- | --- | --- | --- |
| Continuous | $X = (1 - p_x)X^* + p_x\nu_x$ | $p_x = \text{const}$ | $\nu_x \sim \mathcal{N}(0, 1)$ |
| | $Y = (1 - p_y)Y^* + p_y\nu_y$ | $p_y = \text{const}$ | $\nu_y \sim \mathcal{N}(0, 1)$ |
| | $Z = (1 - p_z)Z^* + p_z\nu_z$ | $p_z = \text{const}$ | $\nu_z \sim \mathcal{N}(0, 1)$ |
| Discrete | $X = (1 - \alpha_x)X^* + \alpha_x\nu_x$ | $\alpha_x \sim \text{Ber}(p_x)$ | $\nu_x \sim \text{Ber}(0.5)$ |
| | $Y = (1 - \alpha_y)Y^* + \alpha_y\nu_y$ | $\alpha_y \sim \text{Ber}(p_y)$ | $\nu_y \sim \text{Ber}(0.5)$ |
| | $Z = (1 - \alpha_z)Z^* + \alpha_z\nu_z$ | $\alpha_z \sim \text{Ber}(p_z)$ | $\nu_z \sim \text{Ber}(0.5)$ |

The introduction of impurity in the discrete-variable case is slightly more involved because simple addition of two binary variables does not result in a binary variable. We defined the error terms *ν*_*x*_, *ν*_*y*_, *ν*_*z*_ as standard Bernoulli random variables. We then introduced switching variables *α*_*x*_, *α*_*y*_, *α*_*z*_ modeled by Bernoulli random variables, but this time with varying probability of heads and tails. The observables are obtained by randomly switching between the ground truth variables and the error variables using the switching variables. The probabilities *p*_*x*_, *p*_*y*_ and *p*_*z*_ of the switching variables are the discrete analogue of impurity fractions, as they are equal to the mean values of the switching variables.

In results section we study the performance of the tripartite metrics as function of impurity fractions and data size. To do so, datasets of desired size are sampled from the observable models. Since there are three impurity fractions, one for each of the three observables, we further reduce the number of parameters by designing three different impurity strategies, all of which have only one parameter (table 3). The impurity fractions used in the plots of the main text will refer to this single parameter.

**Table 3:** Three observable models. In the pure sources model (PureSrc), only the target observable *Z* has non-zero impurity, the sources were equal to the underlying ground truth variables. In the impure source *X* model (ImpureX) both the target *Z* and the source *X* observables have impurities (equal impurity fractions), whereas the source *Y* is kept pure. In the impure model (Impure), all three observables have impurities (equal impurity fractions).

| Impurity type | Impurity Fraction |
| --- | --- |
| **PureSrc** | $p_x = 0$ |
| | $p_y = 0$ |
| **ImpureX** | $p_x = p_z$ |
| | $p_y = 0$ |
| **Impure** | $p_x = p_y = p_z$ |

### 2.3 Significance Testing

As a standard method, we employed permutation testing to assess significance of the estimated information atoms. The above-described observable models were used to produce datasets of three observable variables *X*, *Y* and *Z*. Data size of *N*_*sample*_ = 10000 was used everywhere, except when the dependence on data size was investigated. For each dataset, the model information atom was computed. The information atom was then re-computed after permuting the data along the target variable *Z*. This approach is more robust than permuting all three variables, as the metric implementations in practice may be sensitive to source correlations, even though theoretically source correlations should have no impact on the result. This procedure was repeated multiple times (*N*_*test*_ = 10000), obtaining the distributions of the information atom for original and permuted data. The testing threshold corresponding to the desired p-value (0.01) was estimated as the corresponding quantile of the empirical shuffled distribution of the information atoms. The testing threshold was then used to test significance of individual original datapoints, computing the fraction of significant information atoms. If the computed fraction significantly exceed the permutation-test p-value (based on a binomial test, p-value 0.01), we say that the information atom is above shuffle. However, for clarity of presentation, we did not present the value of the binomial test in the main text figures, as the significance of this test is qualitatively evident from the distribution of sample points with respect to the threshold. The testing threshold was independently estimated for all experiments, as it may depend on impurity fractions and data size.

To provide more conservative testing thresholds in view of the bias that we detected for all metrics (see results), we developed an adjusted testing procedure. To produce the conservative testing threshold, the samples were drawn from the corresponding adversarial distribution under the adjusted null hypothesis (see section 3.3), and the corresponding threshold was estimated from the empirical distribution as for the permutation test. The main difference is that the adjusted procedure does not employ data permutation, but directly tests against the worst-case scenario model. Such approaches are a standard way of testing estimators over composite null hypothesis, for example, via a likelihood-ratio test [10]. Similar procedures are commonly used for testing functional connectivity estimators [58].

## 3 Results

We studied the specificity of information atom estimation in simulated ground truth data. We investigated the effect of varying multiple different parameters (see fig. 4A). We tested each of the metrics introduced above (PCorr, VP, and PID) on each of the three ground truth models that were constructed as examples of exactly one underlying information atom: *R*(*X, Y* → *Z*), *U* (*X* → *Z*|*Y*), and *S*(*X, Y* → *Z*), respectively (**ModelR**, **ModelU**, and **ModelS**; see section 2.2.1). If the estimated information atom type matches the type exhibited by the model, we are looking for true positives / false negatives. Otherwise, we are looking for false positives / true negatives. Further, we explored three different observable models (pure source model **PureSrc**, impure source *X* model **ImpureX**, and impure model for both sources **Impure** (see section 2.2.2).

**Figure 4:**
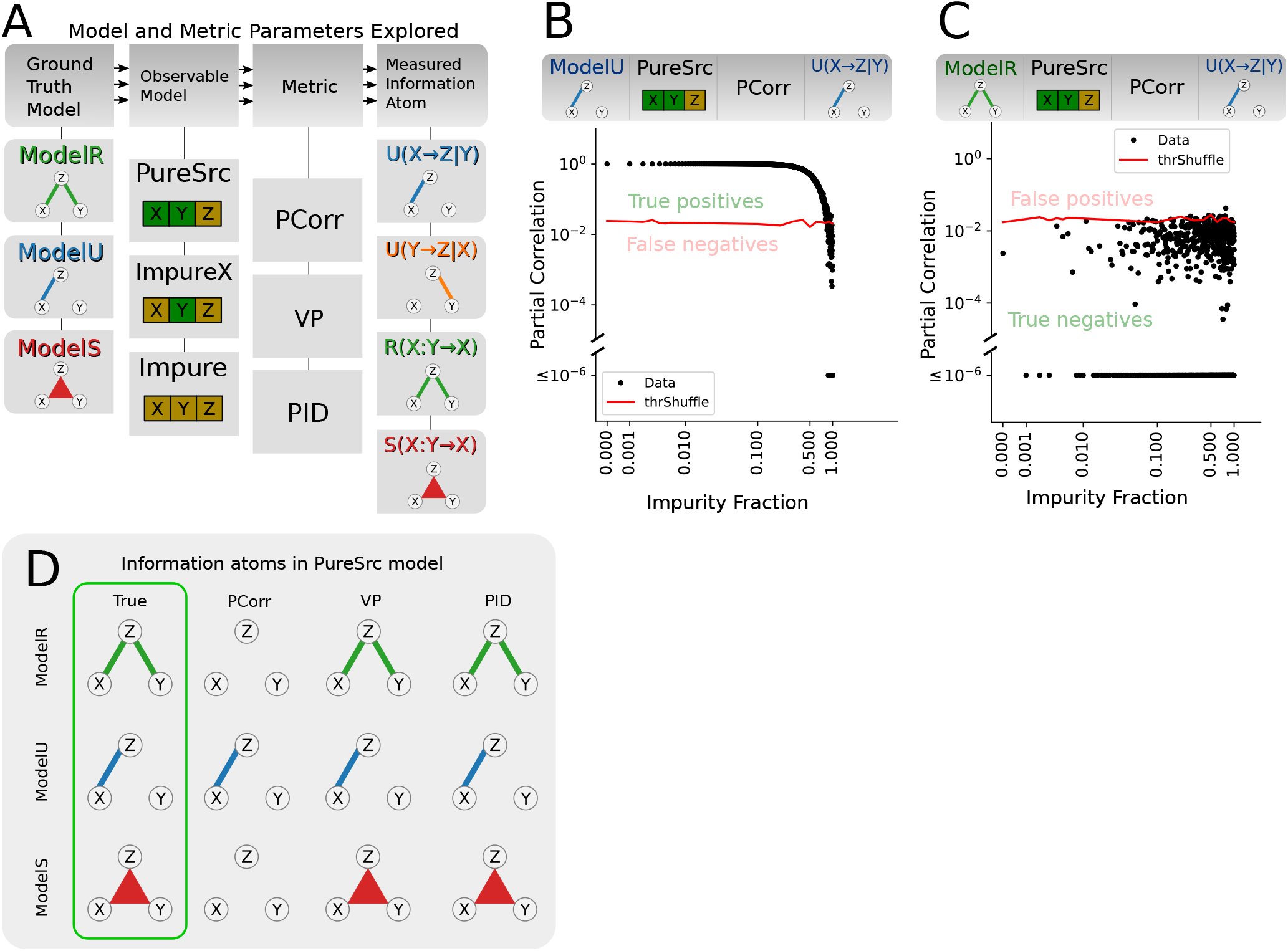
Performance of tripartite analysis metrics on PureSrc model. A) We explored three ground truth models (ModelR, ModelU, ModelS), three observable models (PureSrc, ImpureX, Impure), three metrics (PCorr, VP, PID), which each report 4 different information atoms (except PCorr, which only reports *U* (*X* → *Z*|*Y*) and *U* (*Y* → *Z*|*X*)). In the observational model, green color denotes pure variables (no unexplained variance), and yellow denotes impure variables. B) *U* (*X* → *Z*|*Y*) estimated using PCorr for the pure source ModelU. Plotted is the PCorr magnitude as function of impurity fraction of the model. Red line is the confidence threshold corresponding to p-value of 0.01 for permutation testing. For most impurity fractions the information atom values are significant, correctly resulting in true positives. C) Same as B, but for ModelR. For all impurity fractions, most of the estimated information atom values are not significant, correctly resulting in true negatives. D) Sketch of the detected information atoms for impurity fraction of 0.25 as function of metric (rows) and ground truth model (columns). Line thickness indicates fraction of significant information atoms (permutation-test, p-value 0.01). Emphasized in green are the theoretically expected results for the underlying ground truth model. All metrics correctly identify true positives and true negatives in each model.

First, we showed that the metrics performed as expected in the case of idealised PureSrc observables (impurity fractions *p*_*x*_ = *p*_*y*_ = 0, *p*_*z*_ ≥ 0). We then showed that relaxation of this idealised assumption (*p*_*x*_ ≥ 0, *p*_*y*_ ≥ 0) quickly leads to false positives in all metrics. We exploreed in how far this effect depended on impurity fraction and data size. Finally, to alleviate this problem, we propose to test the information atoms using an adjusted null hypothesis. We performed such tests on simulated data for all metrics. We found that this alternative testing approach helps to eliminate false positives at the expense of increasing false negatives in weaker results. While only selected model and parameter combinations are presented in the main text, all model and parameter combinations are comprehensively shown in the supplementary material. In addition, supplementary material contains results for several validation studies discussed below.

### 3.1 Low false positive rate for pure source variables

First, we asked whether metrics for estimating tripartite functional relations perform as expected in the idealised pure source scenario, i.e., when they have access to the pure (uncorrupted) values of the source variables but receive impure values of the target variable. For each model and metric, we generated distributions of the information atoms for the model data and shuffled data and used the shuffled results to test the significance of the model results (see Methods). We explored the relation between model and shuffle distributions as a function of the target variable impurity fraction. The example *U* (*X* → *Z*|*Y*) values of the PCorr metric for ModelR and ModelU are plotted in fig. 4B and C. All other cases are given in Supplementary Figures 1-14. In fig. 4B we plotted the PCorr *U* (*X* → *Z*|*Y*) for ModelU. For most values of impurity fractions, *U* (*X* → *Z*|*Y*) values for the model data (black) exceeded the permutation testing threshold (red), resulting in true positives. For very large impurity fractions the information atom values did not exceed shuffle, resulting in false negatives, which is expected because the functional relation becomes negligible compared to impurities. In fig. 4C we plotted the PCorr *R*(*X* : *Y* → *Z*) for ModelU. As *R*(*X* : *Y* → *Z*) is not present in ModelU, we expected most of the information atoms estimated from model data not to exceed the testing threshold, which is exactly what we observed.

The summary of all tests is sketched in fig. 4D. We found that all metrics are significant and specific in discriminating between the different models for a broad range of target variable impurity fraction *p*_*z*_. This suggests that all studied metrics are indeed robust and useful at detecting the true underlying information atoms in the case of pure sources.

### 3.2 High false positive rate for impure source variables

Next, we investigated the scenario when the source variables are not pure (observable models ImpureX and Impure; see Methods). In contrast to the PureSrc observable model, both the ImpureX and Impure models resulted in a high fraction of false positives, most prominently for the ModelR ground truth model (fig. 5 E and F). The difference between the ImpureX and Impure observable models was most pronounced in the ModelR. For the ImpureX model, false positives were found only in *U* (*Y* → *Z*|*X*), whereas for the Impure model this was the case for both *U* (*X* → *Z*|*Y*) and *U* (*Y* → *Z*|*X*). This suggests that for the unique information atom estimation, the impurities in the confounding (conditional) variable are significantly more dangerous than those in the target or the primary source. This result was similar for all three metrics, with the fraction of false positives jumping to 100% already for small impurity fraction values. Furthermore, for the PID metric, false positives in *S*(*X* : *Y* → *Z*) also jumped up to 100% for ModelR. Finally, impurities in the main source *X* for ModelU caused the fraction of false positives in *R*(*X* : *Y* → *Z*) to rise up to 50% for the VP and PID metrics. The estimates for ModelS remained robust to impurity in all metrics.

**Figure 5:**
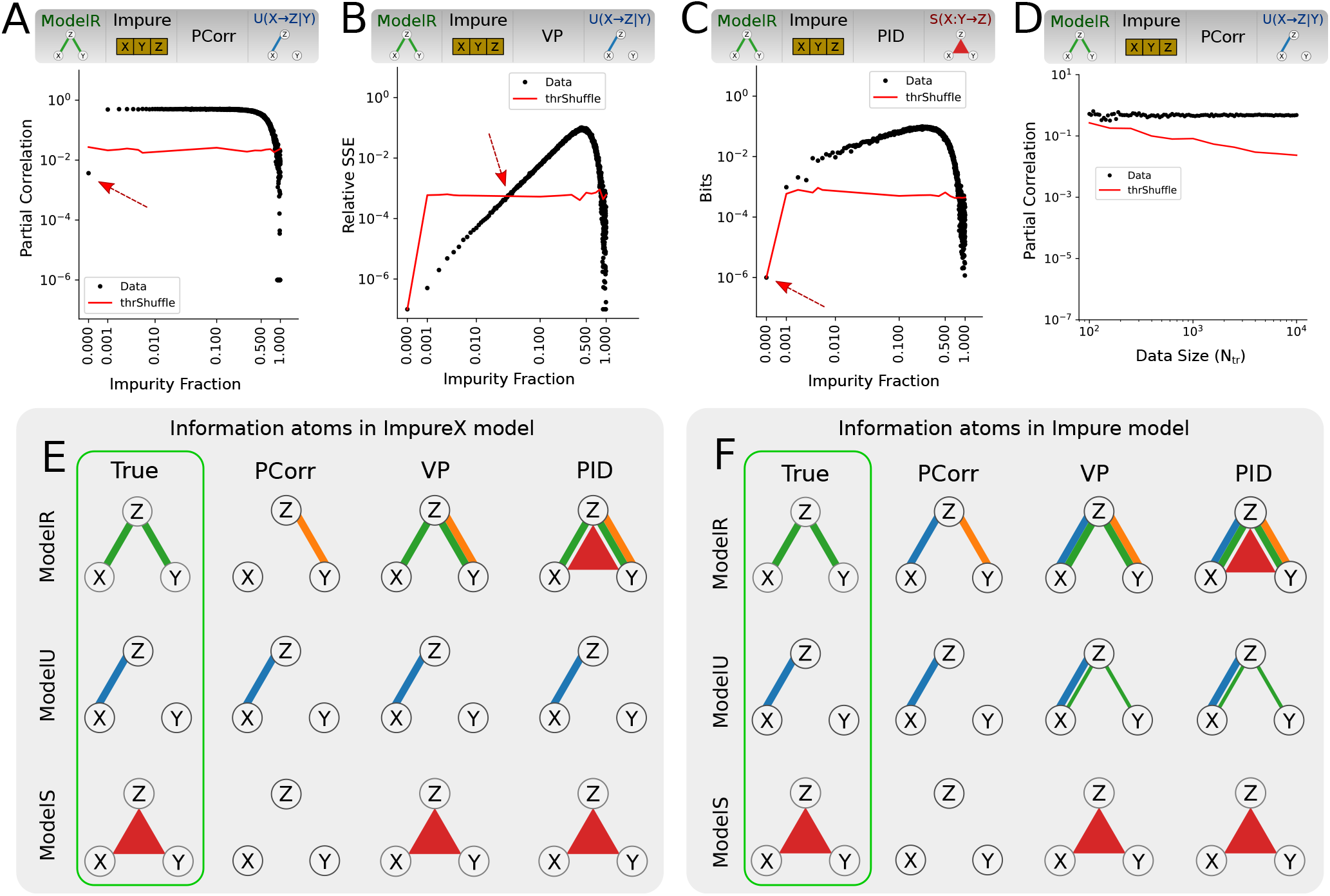
Performance of tripartite analysis metrics on model data with impure source variables. A) PCorr *U* (*X* → *Z*|*Y*) values as function of the impurity fraction for the Impure ModelR. Red line denotes 1% significance threshold based on permutation test (same in B-D). Red dashed arrow indicates transition from true negatives to false positives (same in B, C). B) Same as A, but for VP *U* (*X* → *Z*|*Y*). C) Same as A and B, but for PID *S*(*X, Y* → *Z*). D) PCorr *U* (*X* → *Z*|*Y*) as function of the data size *N*_*tr*_ for a fixed impurity fraction of 0.25 for the Impure ModelR. E) Sketch of the detected information atoms for ImpureX model for impurity fraction of 0.25. Line thickness indicates the fraction of significant information atoms (permutation-test, p-value 0.01).

We thoroughly validated these results. Firstly, we checked the dependence of the results on impurity fraction (fig. 5A, B, C). We found that false positives, such as in Impure ModelR model, jumped up to 100% for low impurity fraction values, and remained at 100% for a broad range of impurity fractions. For PCorr *U* (*X* → *Z*|*Y*) and PID *S*(*X, Y* → *Z*) atoms, impurity fractions of already 0.001 were sufficient to cause false positives. For VP *U* (*X* → *Z*|*Y*) the rise of false positive values was not that fast, requiring values of at least 0.02 to surpass the significance threshold. Importantly, the largest false positive information atom values were comparable with true positive values, suggesting that at least the weaker true positives cannot be discriminated from the false positives based on their magnitude.

Secondly, we checked if the observed false positives were due to insufficient data by studying the asymptotic behaviour of the false positives with increasing data size (fig. 5D, Supplementary Figures 1-14). We found that the effect sizes of the false positive information atoms actually increased with data size, instead of decreasing, suggesting that the false positives were caused by metric bias, not variance. Importantly, information atom values for model data were comparable for different data sizes, whereas the permutation-based testing threshold expectedly decreased with data size, suggesting that the false positives were due to a bias that cannot be fixed with increasing data size.

Finally, the PID metric required additional validation. It is currently designed to work with discrete data and therefore requires data binning to be applicable to continuous-variable data distributions (see Methods). We tested the effect of binning on all models for the Impure model (Supplementary Figure 18). We found that binning can have a dramatic effect on false positives, especially by increasing false positive rate in unique information atoms for ModelR and false positive rate in redundant information atoms for ModelU. While we found the best results with the smallest binning (2-bin discretization), this is likely due to the specific choice of our continuous ground truth models. In general, great care needs to be taken when performing binning. Studies using binning procedures should report the number of bins used, as well as verify if the results hold qualitatively if the bin number is changed. We also checked whether the effects observed using the PID metric could to some extent be caused by the binning procedure itself. To test this we designed a purely discrete version of all of the models (see Methods) and repeated the above validation procedures (Supplementary Figure 17). While there were small quantitative differences, the false positives of the discrete model similarly became significant for very small impurity fractions and remained significant for a broad range of impurity fractions. This suggest that in impure source scenarios the false positive effects are intrinsic to the metric itself.

With this, we conclude that all the considered metrics possess biases in impure source variable scenarios, emerging even for small impurity fractions. Thus, if applied to experimental recordings, permutation-testing of significance for all the considered metrics can be highly misleading.

### 3.3 Adjusted null hypothesis for significance testing of tripartite metrics with improved specificity

To reduce the fraction of impurity-caused false positives in the tripartite metrics, we developed a testing procedure that accounts for biases in the above metrics.

Let *S* be the set of all models for which the true value of the information atom of interest is zero. In this section, when the word ”model” is used alone, we mean the combination of both the ground truth and the observable model.

Let us consider the original permutation test in greater detail. In a permutation test, the test statistic *T* is the information atom value. The null hypothesis *H*_0_ is that the information atom value comes from the distribution that is produced by random permutations of the original data. The aim of the testing procedure is to find a testing threshold Θ. The probability that the information atom value exceeds this threshold given *H*_0_ is called the p-value, and is small (e.g. 0.01).

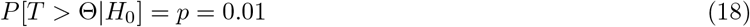

Thus, if the estimated information atom value exceeds this threshold, *H*_0_ is rejected, and it is concluded with confidence level 1 − *p* that the information atom is significant. The main problem with this approach is the choice of *H*_0_. It is implicitly assumed that the permutation-induced distribution of the estimated information atom is representative of that distribution for all models in *S*. As shown in the previous section, this assumption does not hold for the considered tripartite metrics if source variables are impure. The conservative solution designed here is to select the null hypothesis representing the precise scientific question. The adjusted null hypothesis *H*_*adj*_ is that the model that produced the data comes from *S*.

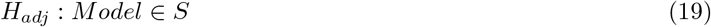

For simplicity, we used the information atom as a test statistic, although more sophisticated test statistics may yield even better results (see discussion). If the estimated information atom value exceeds the testing threshold for *H*_*adj*_ , we may reject all models from *S*. If we are to select only one model *M* ∈ *S* as a null hypothesis, we can obtain the testing threshold Θ_*M*_ for that smaller null hypothesis. The testing threshold Θ_*adv*_ for *H*_*adj*_ is the largest testing threshold over all of the smaller null hypotheses. Thus, the aim is to find a model in *S* which produces the highest possible testing threshold, and use that testing threshold for testing the real data. We will call this model the *adversarial model*.

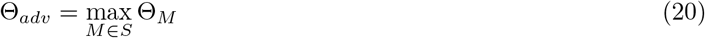

In general, analytic distributions of information atom values are hard to impossible to derive, thus we resort to numerical methods (fig. 6A). Optimization over all possible non-linear models is clearly unfeasible numerically and may require deep theoretical work specific to each metric, which is out of the scope of this study. Instead, we restricted our attention to the same model family that was used to create the data, namely, to linear ground truth models with a quadratic coupling term and to additive impurity observational models. As a further simplification, we only studied corner case adversarial ground truth models, with only a single information atom present at a time. Since for each case we considered only a single ground truth model and a single observational model, we could numerically find the impurity fraction values that produced the highest adversarial information atom values (worst bias).

**Figure 6:**
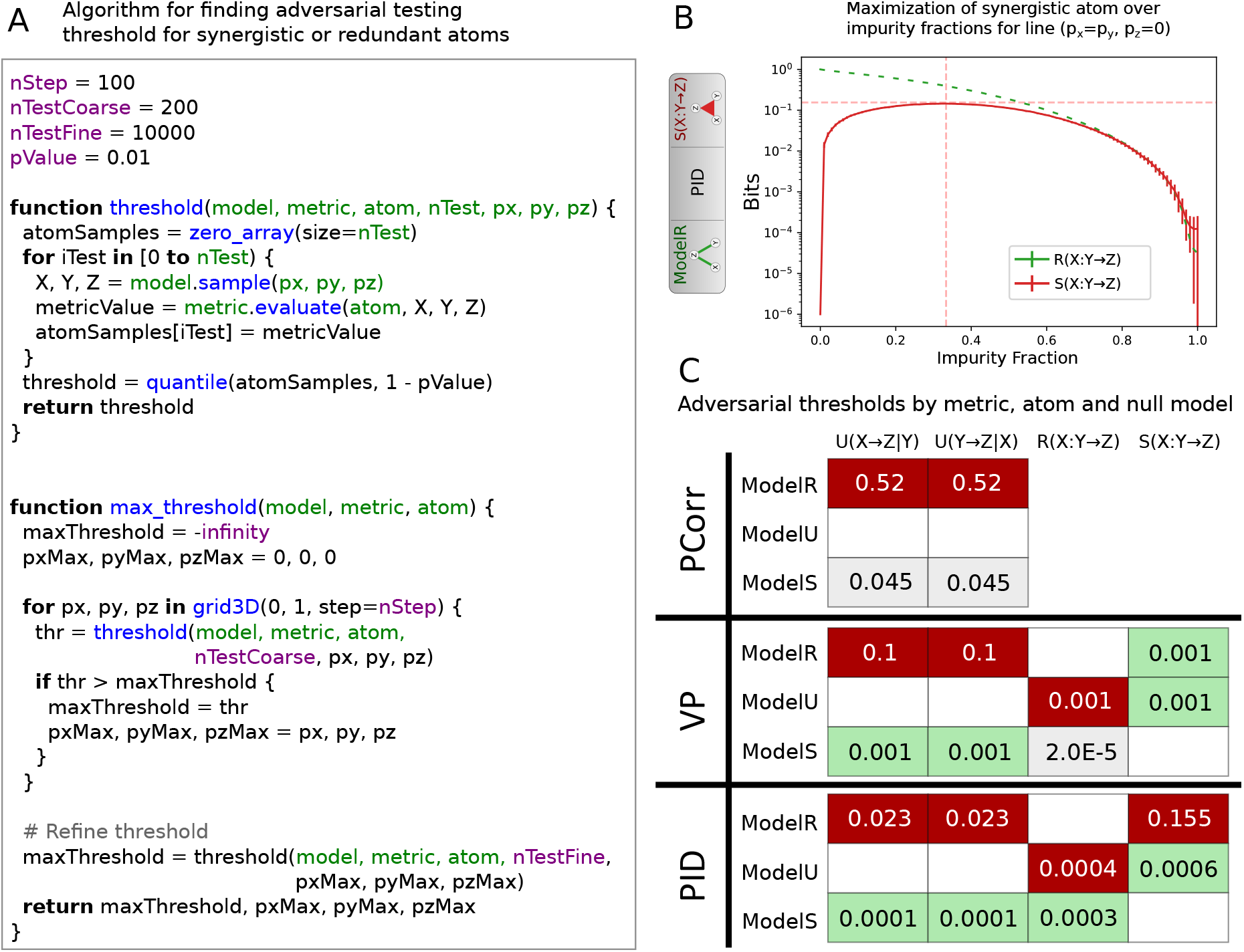
A) Algorithm to determine the adjusted testing threshold for redundant and synergistic information atoms. The function *threshold* finds the testing threshold for a given ground truth and observable model. Function *max threshold* maximizes the testing threshold over all observable models. For unique information atoms, the same algorithm would iterate over a line *p*_*x*_ = *p*_*y*_ = *p*_*z*_ instead of a 3D grid. B) Distribution of false positive *S*(*X* : *Y* → *Z*) (red curve) for PID metric and ModelR model as function of impurity fraction along the line *p*_*x*_ = *p*_*y*_ = *p*_*z*_. Corresponding true positive *R*(*X* : *Y* → *Z*) values (green) are plotted for comparison. Vertical dashed line denotes the impurity fraction with maximal expected false positive *S*(*X* : *Y* → *Z*) value. Horizontal dashed line denotes the 1% upper percentile of *S*(*X* : *Y* → *Z*) at that impurity fraction, corresponding to the p-value 0.01 testing threshold for *H*_*adj*_. C) Estimated testing threshold values for *N*_*tr*_ = 10000 for PCorr, VP and PID metrics. Threshold values correspond to the horizontal dashed line in B) and similar plots (see Supplementary Figures 15, 16). Thresholds are computed for every metric (rows) and every information atom (columns) combination. White (empty) squares indicate true positives. Green squares indicate thresholds that are consistent with permutation test (no false positives). Red squares indicate largest false positive threshold for a given information atom (one per column).

The general strategy is to sweep over ground truth models and observational model parameter values, generate the data from each model to get the empirical distribution, find the testing threshold for each model, and then select the model with the highest testing threshold (the adversarial model). After such parameters are found, the estimated information atom value can be tested for significance against the null model with those parameters. We estimated the information atom distribution under *H*_*adj*_ for each information atom type and each model where that atom is a false positive. For example, for *S*(*X* : *Y* → *Z*), we considered ModelR and ModelU as adversarial models, but not ModelS. For each such distribution, we computed the testing threshold as the upper quantile of the empirical distribution corresponding to the selected p-value (here 0.01).

First, we discuss the unique information atoms, as they are are somewhat different from the other two information atoms. In principle, the unique information atom in the ModelR can become arbitrarily prominent if the impurity fraction in one of the redundant source variables is arbitrarily large. In such situations, the true information atom value is impossible to estimate unambiguously (see discussion). Instead, we addressed a sub-problem in which all variables have the same impurity fractions (*p*_*x*_ = *p*_*y*_ = *p*_*z*_), in other words, using the Impure model as the adversarial model. This situation can emerge in neuroscience. For example, recordings of multiple neuronal variables may be corrupted by observational noise of the same distribution. Conceptually, the unique information atoms emerge here as false positive because impurity corrupts the two redundant source variables in a different way, making them individually significant as predictors of the target variable. This collaborative effect between two impure sources is great for improving prediction accuracy of the target, but is certainly undesirable as an estimator of unique information atom significance. We found the maximum likelihood estimate for impurity fractions that produced the highest expected information atom value for false positive unique information atoms via a grid search. Impurity fraction values between 0 and 1 were split into 100 steps, then for each step the information atom value was resampled 200 times, computing the expected value and the 1% upper percentile threshold value. Once the impurity fraction resulting in highest threshold value was found, the model was resampled 10000 times for that impurity fraction, finding a more precise estimate of the threshold. We refer to this value as the *adjusted testing threshold* for unique information atoms. The distribution of the false positive information atom value was found to change smoothly with impurity fraction, suggesting that the loss from using an overly conservative threshold is minimal for a large range of impurity fraction values (fig. 6, B). This procedure was repeated for all metrics. Further, we explored data sizes in the range of 1000-10000 data points, and found that the threshold was almost constant for data size values within this range, with values about 3-4% higher for lower data sizes (Supplementary Figures 15,16).

Second, we aimed to correct the bias in redundant and synergistic information atoms. Unlike unique information atoms, false positive synergistic and redundant information atoms did not exhibit unbounded growth with impurity fraction asymmetry between source variables. Hence, it was necessary to find the maximum likelihood solution over all combinations of all three impurity fraction parameters. We used a grid search with a coarse grid of 10 steps, discretizing the impurity fraction values of each variable between 0 and 1. For each set of parameter values, we performed 400 tests and estimated the mean information atom value. The grid search revealed that the highest information atom values could be observed when the target variable impurity fraction was very low, but the impurity fractions of both the source variables were equal. This result held for all data sizes considered, and was further confirmed by a finer grid search in 2D, holding the target variable impurity fraction fixed. We proceeded to find the 1% threshold using the same procedure as for the unique information atom. Similarly to the unique information atom, the MLE synergistic and redundant information atom distributions were relatively smooth with respect to impurity fraction (fig. 6B), and consequently changed little with data size, with larger values for smaller data sizes. We summarized the testing thresholds for *N*_*tr*_ = 10000 for all considered metrics, information atoms and adversarial models in fig. 6C.

We then used the obtained thresholds to re-test data from all metrics and models (fig. 7F). We found that our procedure eliminated false positives in all considered metrics and models in Impure model. Results were qualitatively similar in ImpureX model, with the exception of *U* (*Y* → *Z*|*X*) atoms, where the false positives remained for the reasons discussed above. Here we present the detailed analysis for a selection of metrics and information atom types (fig. 7A, B, C, D) for the Impure model. The plots are the same as the previously shown plots (fig. 5A, B, C, D) respectively, except there is an additional threshold line (purple) denoting the adjusted testing threshold. All other model and information atom combinations are presented in Supplementary Figures 1,3,5,7,9,11,13. As a limitation of our approach, stricter thresholds also resulted in an increase of false negatives. For example, false negatives in PCorr *U* (*X* → *Z*|*Y*) for ModelR only appeared for impurity fractions above 0.8 when using permutation-testing, but started appearing already for impurity fraction of 0.5 when using the adjusted testing procedure (fig. 7A). Qualitative behaviour was the same for all true positives when tested against *H*_*adj*_, but transition impurity fractions varied (Supplementary Figures 1,3,5,7,9,11,13). Further, we inspected the adjusted testing procedure as a function of data size. We plotted PCorr *U* (*X* → *Z*|*Y*) for ModelR using impurity fraction of 0.25 in (fig. 7E), all other parameter combinations in Supplementary Figures 2,4,6,8,10,12,14. Adjusted testing threshold (purple) changed marginally with data size, decreasing for larger values.

**Figure 7:**
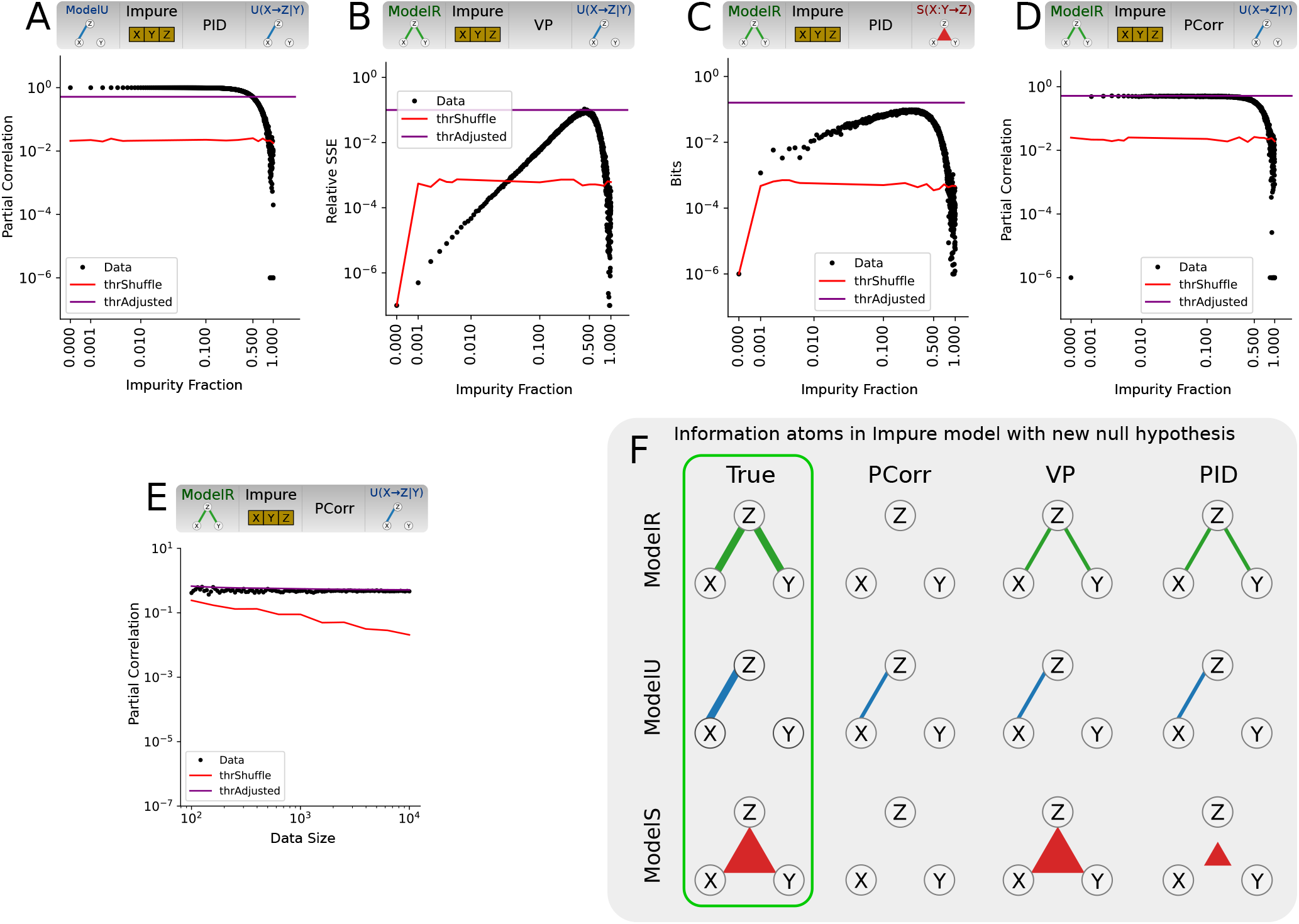
Performance of tripartite analysis metrics on model data with impure source variables, tested against *H*_*adj*_. All plots are for the Impure model. A) PCorr *U* (*X* → *Z*|*Y*) for ModelU, as function of impurity fraction (true positives). Red and purple lines correspond to the permutation testing and adjusted testing thresholds respectively (same for B-E). B,C,D) False positives in PCorr, VP and PID as a function of impurity fraction. Same plots as (fig. 5A, B, C) respectively. E) False positives in PCorr as function of data size for impurity fraction of 0.25. Same plot as (fig. 5D). F) Sketches of the detected information atoms for Impure model for impurity fraction of 0.25. Line thickness indicates the fraction of significant information atoms (tested against *H*_*adj*_, p-value 0.01)

**Figure 8:**
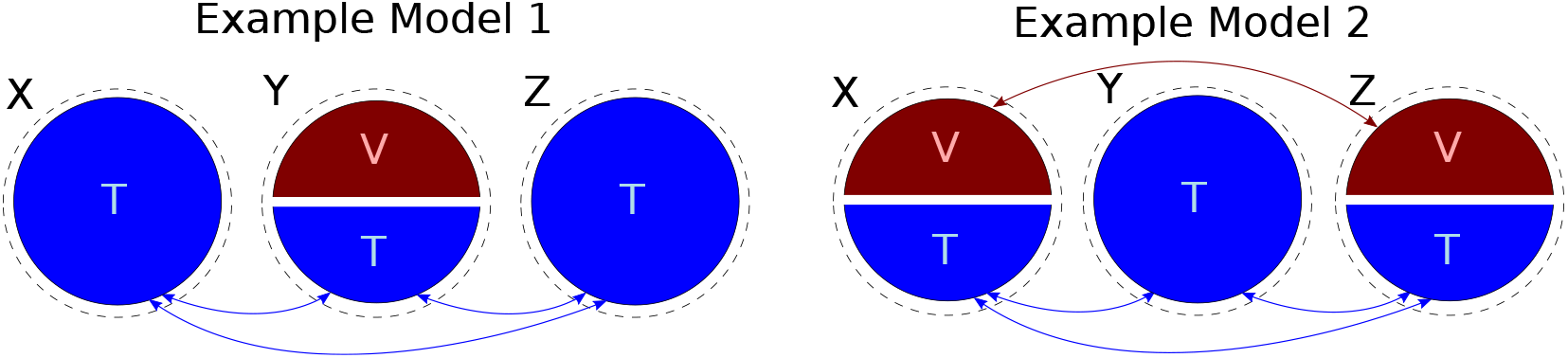
Two different ground truth designs that can produce indistinguishable data. Three populations *X*, *Y* and *Z* redundantly encode a latent variable *T* . In model 1, the population *Y* additionally encodes another latent variable *V* , whereas in model 2 the second latent variable is additionally encoded by *X* and *Z*.

## 4 Discussion

In this work, we studied whether permutation-testing of tripartite functional relation estimators is a robust approach for estimating ground truth relations from simulated data in the presence of data impurities. Three metrics commonly used for such analysis proved significant and specific in the absence of source impurities in the model. However, additions of even small impurities to the source signals resulted in dramatic loss of specificity in all metrics considered, producing up to 100% false positives. We also demonstrated that false positives become even more significant with increasing data sizes, concluding that this problem cannot be fixed by acquiring more data. As a consequence, if applied to experimental data, permutation testing of these metrics could result in falsely detecting pairwise-specific functional connections in a purely redundant system, which is undesirable and misleading. To address this problem, we designed an alternate testing procedure that accounts for model biases in the presence of impurities. Compared to permutation-testing, our conservative test consistently eliminated false positives in the studied metrics, albeit at the expense of introducing more false negatives with increasing impurity fraction. This testing procedure is applicable to any tripartite metric estimating information atoms or related quantities.

Researchers are invited to run the simulations in the provided python code (for a given metric and data size) to find the corresponding conservative testing thresholds that then can be applied to experimental data.

It is interesting to analyse why the false positives highlighted in this work emerge. Importantly, some of the false positives are not due to shortcomings of individual metrics, but rather due to a fundamental ambiguity in data recorded from under-controlled and/or noisy complex systems. For example, consider two following situations (fig. 8). In the first situation, population-average observables *X*, *Y* and *Z* redundantly encode some latent variable *T* . Further, *Y* averages over two different populations of neurons - one that is redundant with *X* and *Z* (encoding *T*), the other unrelated to *X* or *Z* (called *V*). The constant *α* ∈ [0, 1] determines the relative signal strength of the two neuronal populations in *Y* .

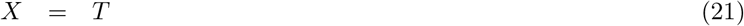

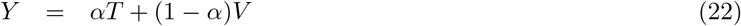

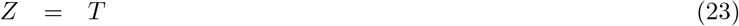

For *α* between (0, 1) (e.g. *α* = 0.5), redundancy is partially destroyed due to averaging over the two populations in *Y* . In this situation, our analysis will find a non-negligible *R*(*X, Y* → *Z*), as well as a non-negligible *U* (*X* → *Z*|*Y*). In the second situation, both *X* and *Z* are averages over two populations of neurons, whereas the population of neurons in *Y* is uniform. The first population in *X* is redundant to the first population in *Z* and to the only population in *Y* (given by the latent variable *T* , same as above). The second population in *X* will be correlated to the second population in *Z*, but unrelated to *Y* (given by the latent variable *V*). Here, the constant *β* ∈ [0, 1] will determine the relative strength of the two neuronal populations in both *X* and *Y* .

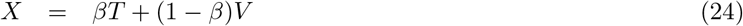

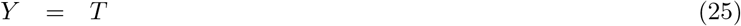

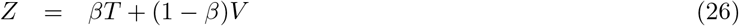

For appropriate values of the constants, the data distribution sampled from the second model can be statistically indistinguishable from the one sampled from the first model. The difference, however, is that in the second situation both *U* (*X* → *Z*|*Y*) and *R*(*X, Y* → *Z*) meaningfully relate to the underlying neuronal interactions, whereas in the first situation *U* (*X* → *Z*|*Y*) may be misleading, since *X* and *Z* do not share a stronger connection than for example *X* and *Y* . To summarize, this example shows that redundant and unique information atoms can become indistinguishable in cases where the additive impurities have different magnitude in *X*, *Y* and *Z*. We recommend to take this fact into consideration for future experimental design and interpretation.

Next, we discuss related research in Functional Connectivity (FC) [33] and Effective Connectivity (EC) [39], and highlight potential implications of our results on estimation of related metrics. Metrics of FC and EC aim to estimate a matrix of pairwise connections between variables (also known as functional connectome [29]), to test if individual connections are significant, to describe the connectivity matrix by means of integral metrics of network neuroscience [7], and to study changes in network connectivity associated for example with learning [6, 73, 74] or disease [14]. Redundancy is a well-known problem in this field as well. Bayesian approaches [32] model the posterior distribution of all parameter value combinations and typically bypass the redundancy problem by comparing the relative evidence of a few biologically-motivated parameter combinations [61]. A Frequentist approach to address the problem is to introduce a strict additional criterion on the specificity of inferred connectivity (such as optimal information transfer [49]) and to iteratively prune connections according to such criterion. Comparison of pairwise and pruned connectivity matrices can be used to approximate the range of possible functional networks [74].

We conjecture that source impurities can negatively affect estimates of time-directed functional connectivity metrics, such as transfer entropy. Such metrics estimate functional connectivity between a past time point of the source signal and the current time point of the target signal, conditioned on the past time point of the target signal. It relies on the metrics similar to those studied here (partial correlation and conditional mutual information) and thus will likely be subject to false positives in the presence of impurity. More precisely, a frequent application is the estimation of transfer entropy between two autocorrelated signals that are also correlated at zero lag. The user may be interested in checking if there is significant functional connectivity at small but non-zero lag, independently from the apparent zero-lag functional connectivity. In this case, the activity values of the past of the source, the past of the target and the current time point of the target will be redundant, and we expect the metric to find no significant lagged functional connections, which may not be the case in the presence of impurity. Nevertheless, the worst-case scenario for transfer entropy is less dire than that for a general tripartite metric. As past and present of the target come from the same signal, the impurity fractions of both of these variables in real data are equal or almost equal, significantly reducing the possible magnitude of false positive unique information atoms. In another study [74], we validated the performance of transfer entropy in the presence of noise for simulated neuronal recordings. We found that the metric was able to correctly reject false positives within a range of low impurity fractions.

During the last two decades, evidence has accumulated in support of the presence of higher-order interactions (tripartite and above) in neuronal populations, including in vivo and in vitro experiments, as well as simulations (see [82] for a review). Most of the analyses were performed in the framework of either information geometry [3] or Maximum Entropy [67, 70]. Both frameworks require fitting the data to a multivariate probability distribution from an exponential family. Comparison of models of different complexity (e.g. via maximum likelihood) is used to determine whether the more complex models involving higher-order terms are better at explaining the observed data. While we did not explicitly investigate the effects of impurity on these frameworks, our current results suggest that these frameworks could be vulnerable to impurity, similar to the simpler models studied in our work. Further, synergy and redundancy have been extensively studied in neuroscience by means of the predecessor of PID, namely the Interaction Information (II) and similar metrics (see [76] for review). Very recently, this metric has been used to demonstrate synergistic encoding of spatial position by neuron-astrocyte pairs [21]. Since II is strongly related to PID and also does not explicitly correct for impurities, we would expect the impurity induced false positives to be just as relevant.

Despite focusing this paper on functional relations between triplets of neuronal signals, our statistical results are general and can see applications outside the scope of neuroscience. Studies of confounding effects, especially by means of partial correlation or partial r-squared are common in econometrics [47, 77], medicine [15], genetics [34, 65], neurochemistry [5], psychology [26, 79] and many other fields. Synergistic effects, among others, have been studied in physical systems [8], ecology [54] and sociology [17]. Further, earlier in this work we provided an example application where all three variables were of neuronal origin. This choice is purely an interpretation of our statistical results and is done for clarity of presentation purposes. All of our findings are equally applicable to scenarios where all or some of the source and/or target variables non-neuronal, such as behavioural or sensory variables. For example

- Functional/Effective connectivity between neurons may be investigated as function of an exogenous variable (e.g. treatment, stimulus or behaviour) in a mixed behavioural-neuronal experiment with one exogenous source.
- Multi-sensory integration in a cortical or sub-cortical brain area [24] could be studied as function of auditory and visual stimuli in a mixed behavioural-neuronal experiment with two exogenous sources.
- The performance of a participant may be analyzed as function of learning time and reward size in a purely behavioural experiment.

Our results rely on several simplifying assumptions, some of which are worth to be improved upon in future studies. First, we computed information atoms using data distribution across trials for a fixed time step. A related question is the study of information atoms across time, for example, in long recordings of resting-state activity. Compared to the former, across-time analysis is complicated by autocorrelation in data. We refer the reader to related recent work addressing autocorrelation effects in functional connectivity estimation [42, 20]. Second, we described estimating information atoms using simultaneous source and target data (zero lag). The tripartite metrics can be estimated with source signals lagged compared to the target, yielding time-directed information atom estimates [78]. Zero-lag estimates can also be thought of as time directed, under the assumption that timescale of signal propagation in the system is faster than a single time step. Importantly, our results apply equivalently to any choice of lag, as selection of arbitrary lags would still result in a three variable empirical distribution. For further reference on interpretation of lagged estimators see [78]. Third, we used a linear model with a quadratic coupling term and a Gaussian additive impurity term. It will be interesting to verify if our results hold for more complex nonlinear ground truth models, non-additive (e.g. multiplicative) impurity and non-Gaussian (e.g. log-normal) impurity distribution. Fourth, our testing procedure relies on several assumptions and simplifications. We assume that false positives are worse than false negatives in exploratory neuroscientific research, since a false detection of a functional relation presumably is more misleading than missing a weaker real relation. Our testing procedure can be made more robust by considering other potential adversarial models, such as non-linear models of higher order, or quadratic models with mixed terms. Sensitivity of our testing procedure can also be improved, reducing the number of false negatives while preserving sensitivity. This is due to the observation that not all of the combinations of information atoms are possible. For example, the maximal value of the false positive *S*(*X* : *Y* → *Z*) for PID and ModelR depends on the true value of the *R*(*X* : *Y* → *Z*), as seen in fig. 6B. Instead of testing one information atom at a time, it may be possible to take advantage of the multivariate distribution of all information atoms simultaneously. Finally, application of our validation approach to more advanced metrics, such as higher-order decompositions [80], continuous information-theoretic estimators [60, 66] and symmetric information-theoretic estimators [63] should provide insight into practical advantages and challenges of these metrics in application to impure neuronal data.

In this work, we presented several applications of tripartite metrics to simulated data, and demonstrated their usefulness in inferring more advanced network features than those provided by pairwise functional connectivity estimators. We conclude that statistical concerns of testing such metrics can mostly be resolved, and hence recommend the use of such metrics in future experimental and computational literature. Further, our work presents an example of how permutation-testing of a novel metric can produce misleading results. Given the popularity of permutation-testing in neuroscience, we recommend extensive theoretical and numerical validation of novel metrics prior to use on experimental data.

## 5 Author Contributions

A.F. constructed proof of principle. Y.S. outlined the coarse manuscript structure. A.F. designed and implemented all simulations, and wrote the manuscript. F.H. and Y.S. reviewed the manuscript.

## 6 Acknowledgements

We thank Joseph Lizier, Patricia Wollstadt and Leonardo Novelli for initial support in using the library IDTxl. We are grateful to Michael Wibral and Abdullah Makkeh for extensive support on theory underlying partial information decomposition, especially in terms of interpretation of results. We thank Peter Rupprecht, Adrian Hoffmann, Christopher Lewis, and many other members of Helmchen Lab for suggestions on improving the manuscript. Finally, we thank Willliam Huber, Ruben van Bergen and Frank Harrell for useful suggestions with respect to our questions on the state of the art in statistical analysis.

This work was supported by grants to F.H. from the Swiss National Science Foundation (No. 310030B 170269) and the European Research Council (ERC Advanced Grant, project No. 670757, BRAINCOMPATH).

## 7 Code Availability

All code used for this project is available in the open source github repository.

https://github.com/aleksejs-fomins/pub-2020-exploratory-analysis/tree/master/analysis-null

Note that this project makes extensive use of another library for general purpose multivariate statistical analysis in neuroscience, developed by the authors during this project.

https://github.com/HelmchenLabSoftware/mesostat-dev

