## Supplementary Material for "Conservative Significance Testing of Tripartite Interactions in Multivariate Neural Data"

February 7, 2022

### 1 Proofs

#### 1.1 Derivation of the Variance Partitioning Redundancy Atom

While it is possible to construct the redundant atom directly from the quadratic model used in the main text, we found it more numerically robust to use the linear model, ignoring the quadratic term. Consider the following 3 linear models

$$Z_{lin,x}(X) = aX \tag{1}$$
$$Z_{lin,y}(Y) = bY \tag{2}$$
$$Z_{lin,x,y}(X,Y) = aX + bY \tag{3}$$

The sum of squared errors for each of the above models can be written in terms of the information atoms. First, we use the full linear model $Z_{lin,x,y}(X,Y)$ to decompose the total sum of squares $SST_Z$.

$$SST_Z = SSE_{lin,x,y} + SSR_{lin,x,y} \tag{4}$$

The sum of squares explained by this model is the sum of squares explained uniquely by $X$, plus the sum of squares explained uniquely by $Y$, plus the sum of squares explained redundantly by $X$ and $Y$ (redundant atom $R(X:Y:Z)$).

$$SSE_{lin,x,y} = U(X \to Z|Y) + U(Y \to Z|X) + R(X:Y \to Z) + SSR_{lin,x,y} \tag{5}$$

Secondly, we can decompose the total sum of squares using the other two models

$$SST_Z = SSE_{lin,x} + SSR_{lin,x} \tag{6}$$
$$SST_Z = SSE_{lin,y} + SSR_{lin,y} \tag{7}$$

As an aside, note that while the explained and residual sum of squares depend on the model function, the total sum of squares $SST_Z$ is only a property of the target data.

$$SST_Z = \sum_i |z_i|^2 \tag{8}$$

The explained sums of squares will contain the redundant atom and the unique atom corresponding to the parameter variable, but will not contain the unique part not present among the parameters

$$SSE_{lin,x} = U(X \to Z|Y) + R(X:Y \to Z) + SSR_{lin,x,y} \tag{9}$$
$$SSE_{lin,y} = U(Y \to Z|X) + R(X:Y \to Z) + SSR_{lin,x,y} \tag{10}$$

Subtracting eq. (6) and eq. (7) from eq. (4), and solving for the redundant atom yields the following definition

$$R(X:Y \to Z) = SST_Z - (SSR_{lin,x} + SSR_{lin,y} - SSR_{lin,x,y}) \tag{11}$$

### 2 Figures

Note that the magnitudes of all information atoms in all log-scale plots are cropped to the minimum value of $10^{-6}$ or $10^{-7}$, as can be inferred from the individual plots. The purpose of this cropping is to focus the logarithmic plot on the more important part of larger information atoms, as well as to avoid over-interpretation of numerical noise.

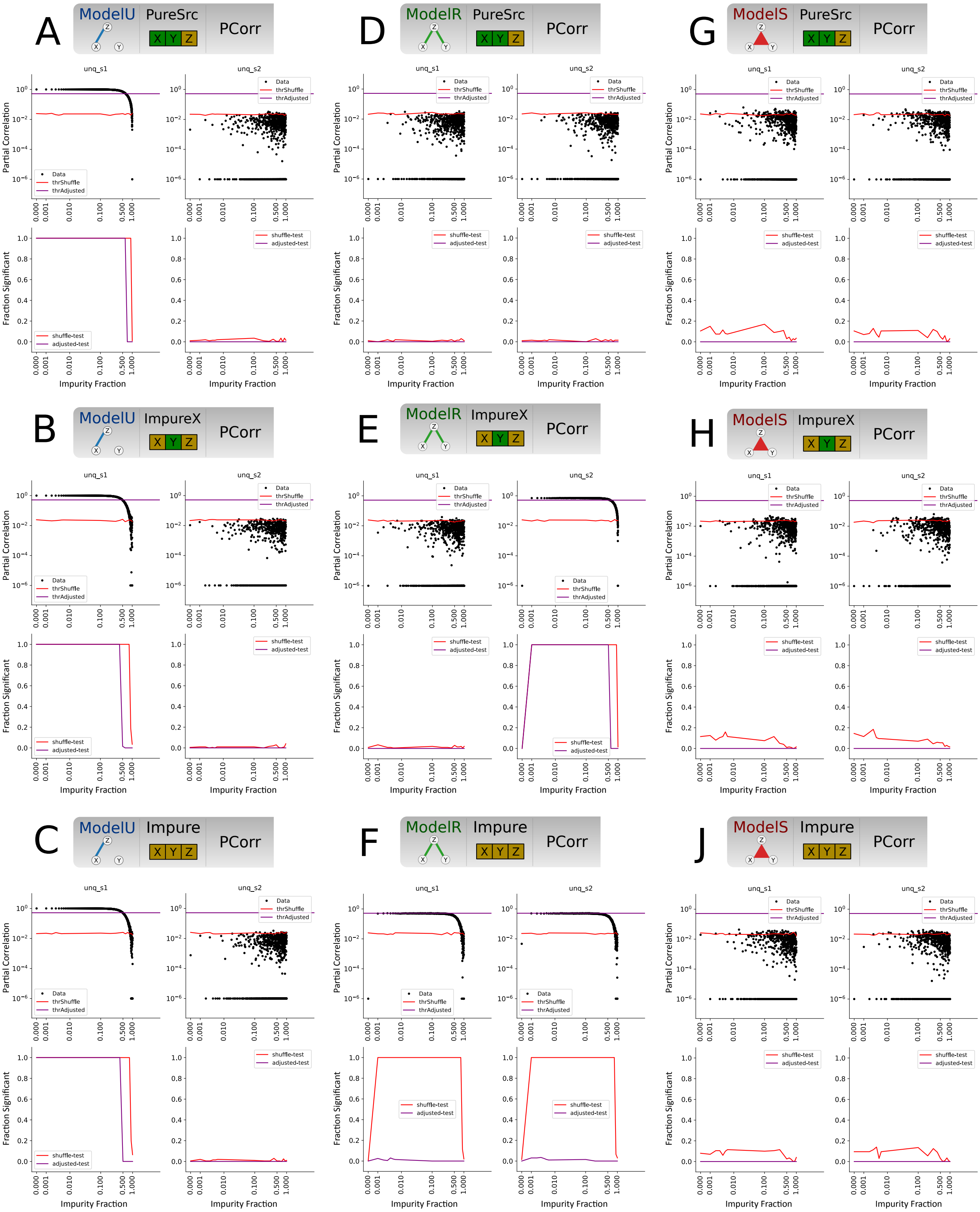


Figure 1: Partial Correlation magnitude (top) and fraction of significant values (bottom) for different ground truth and observable models, as function of impurity fraction for $N_{tr} = 10000$. Red line denotes permutation testing threshold, and corresponding fraction of significant values. Purple line denotes the same for the adjusted conservative test. Columns in each figure denote two unique information atoms $U(X \to Z|Y)$ and $U(Y \to Z|X)$ respectively.

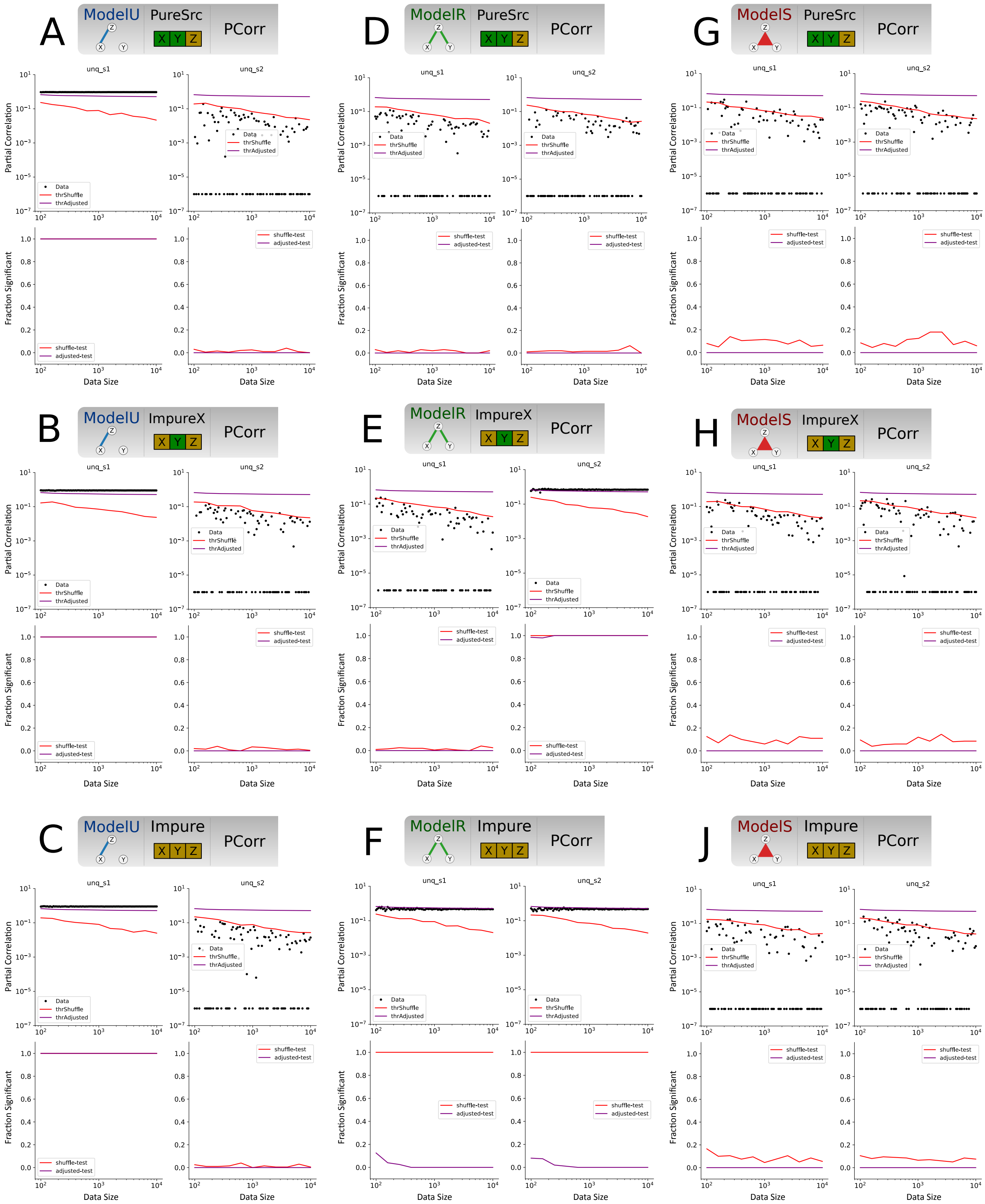


Figure 2: Partial Correlation magnitude (top) and fraction of significant values (bottom) for different ground truth and observable models, as function of data size for fixed impurity fraction 0.25. Red line denotes permutation testing threshold, and corresponding fraction of significant values. Purple line denotes the same for the adjusted conservative test. Columns in each figure denote two unique information atoms $U(X \to Z|Y)$ and $U(Y \to Z|X)$ respectively.

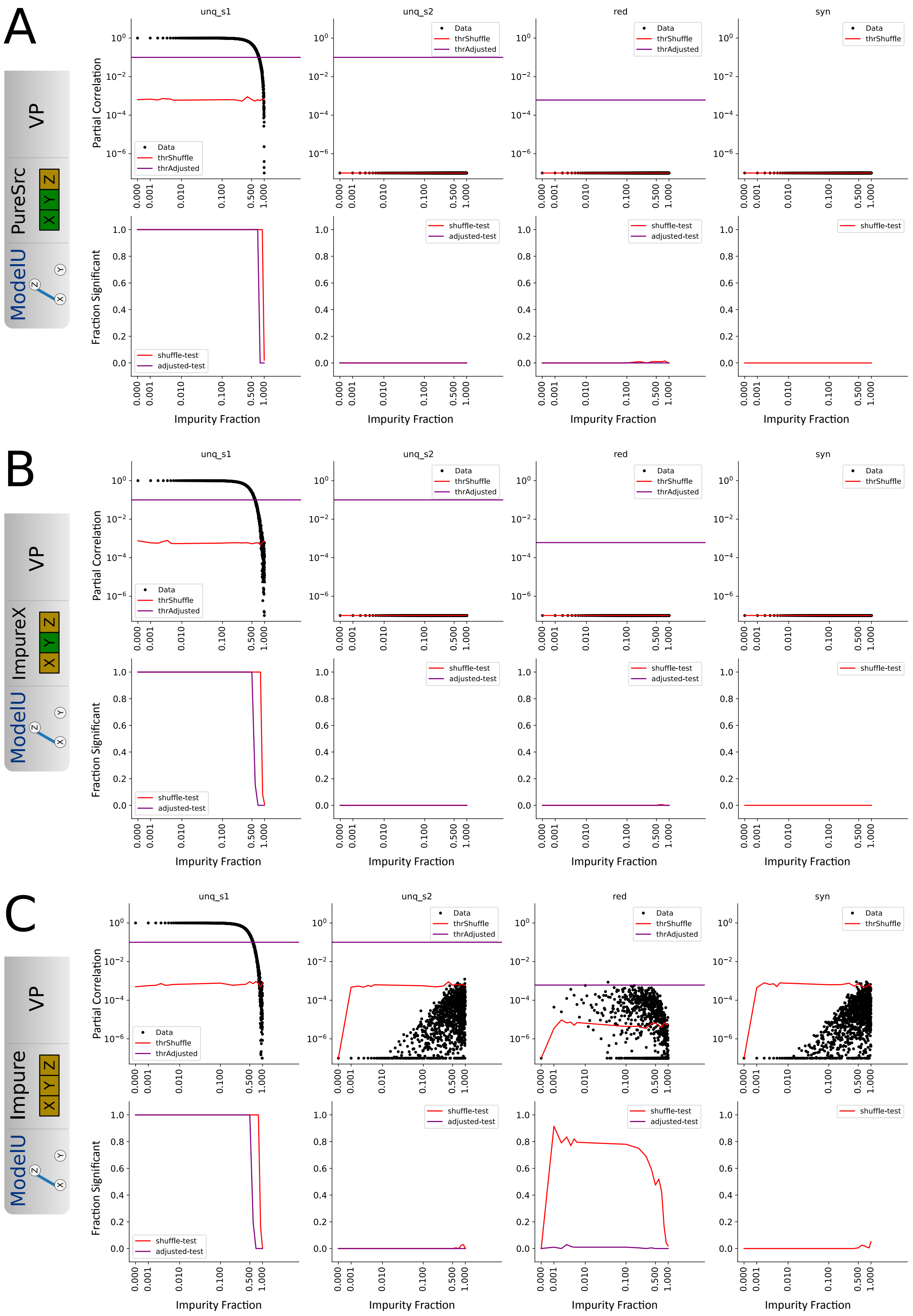


Figure 3: Variance Partitioning magnitude (top) and fraction of significant values (bottom) for ModelU and different observable models, as function of impurity fraction for $N_{tr} = 10000$. Red line denotes permutation testing threshold, and corresponding fraction of significant values. Purple line denotes the same for the adjusted conservative test. Columns in each figure denote information atoms $U(X \to Z|Y)$ and $U(Y \to Z|X)$, $R(X, Y \to Z)$ and $S(X, Y \to Z)$ respectively.

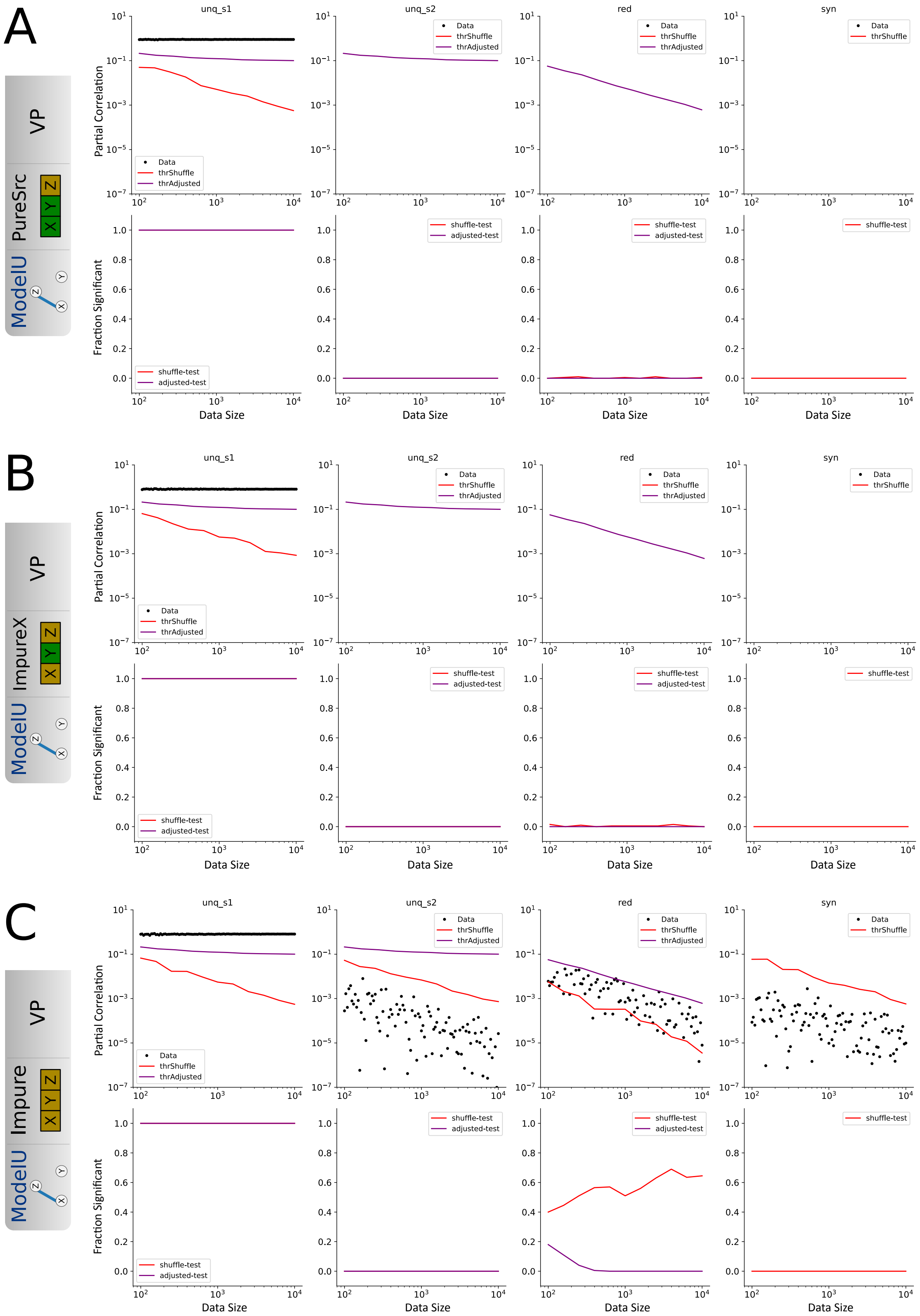


Figure 4: Variance Partitioning magnitude (top) and fraction of significant values (bottom) for ModelU and different observable models, as function of data size for fixed impurity fraction 0.25. Red line denotes permutation testing threshold, and corresponding fraction of significant values. Purple line denotes the same for the adjusted conservative test. Columns in each figure denote information atoms $U(X \to Z|Y)$ and $U(Y \to Z|X)$, $R(X, Y \to Z)$ and $S(X, Y \to Z)$ respectively.

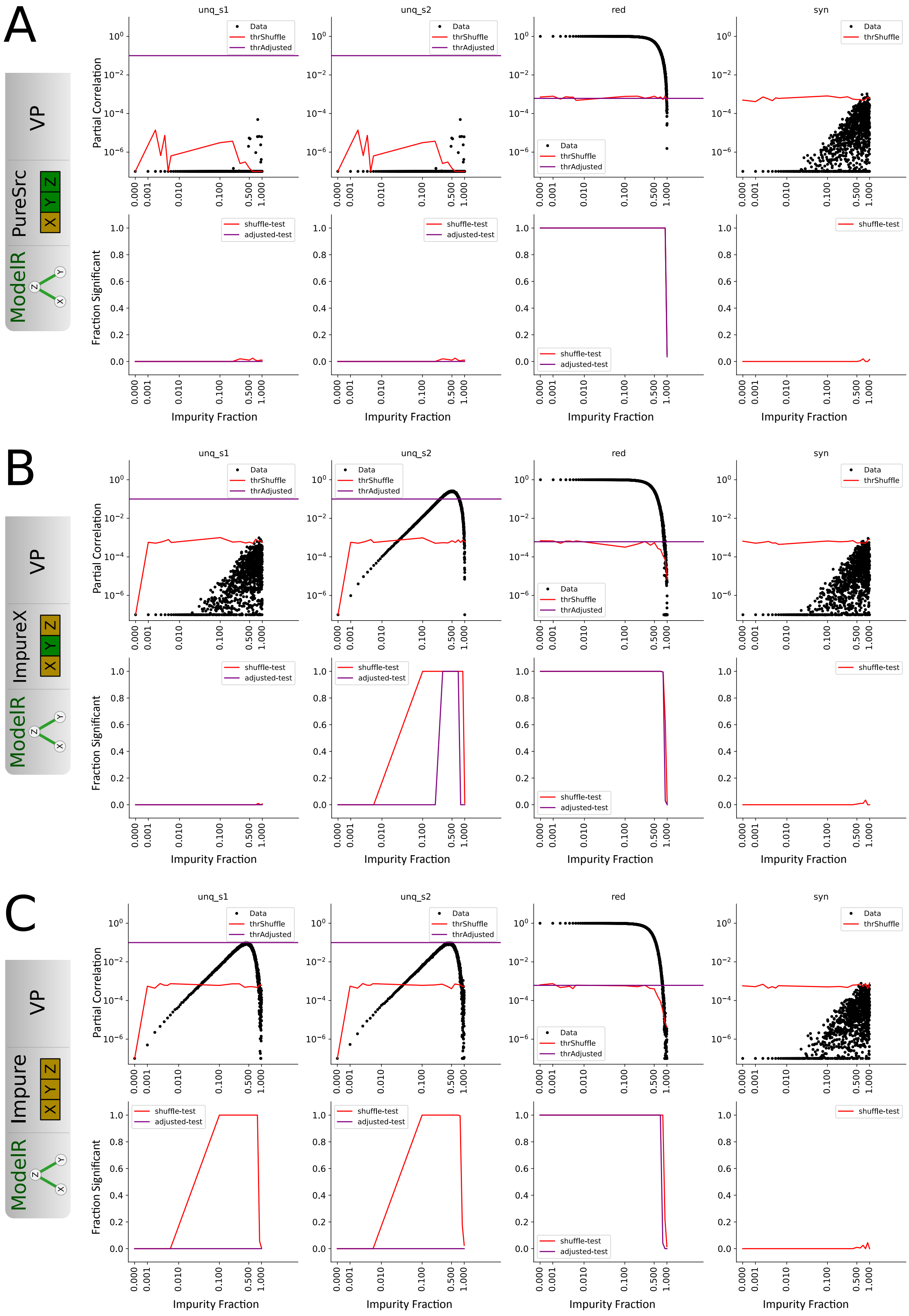


Figure 5: Variance Partitioning magnitude (top) and fraction of significant values (bottom) for ModelR and different observable models, as function of impurity fraction for $N_{tr} = 10000$. Red line denotes permutation testing threshold, and corresponding fraction of significant values. Purple line denotes the same for the adjusted conservative test. Columns in each figure denote information atoms $U(X \to Z|Y)$ and $U(Y \to Z|X)$, $R(X, Y \to Z)$ and $S(X, Y \to Z)$ respectively.

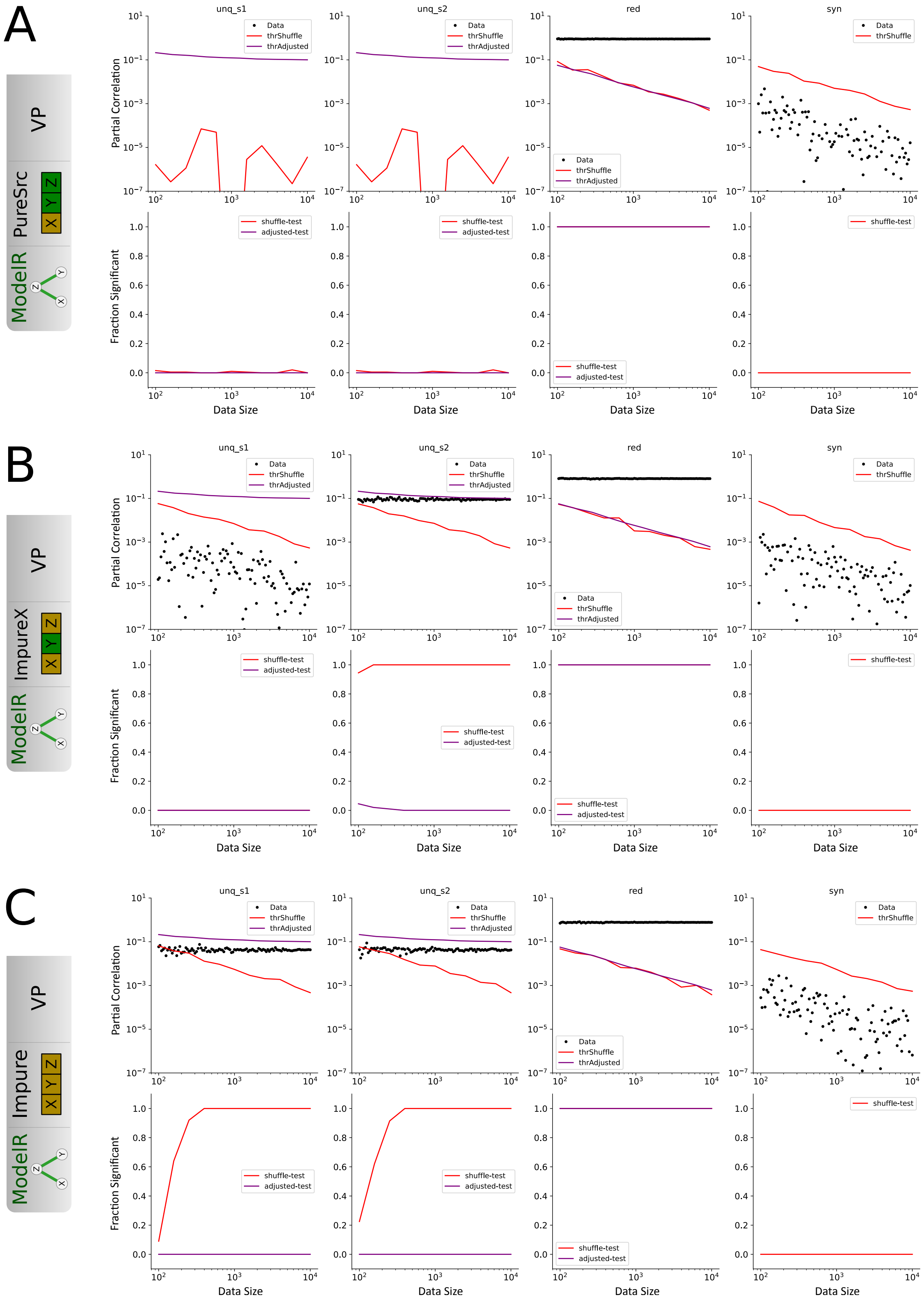


Figure 6: Variance Partitioning magnitude (top) and fraction of significant values (bottom) for ModelR and different observable models, as function of data size for fixed impurity fraction 0.25. Red line denotes permutation testing threshold, and corresponding fraction of significant values. Purple line denotes the same for the adjusted conservative test. Columns in each figure denote information atoms $U(X \to Z|Y)$ and $U(Y \to Z|X)$, $R(X, Y \to Z)$ and $S(X, Y \to Z)$ respectively.

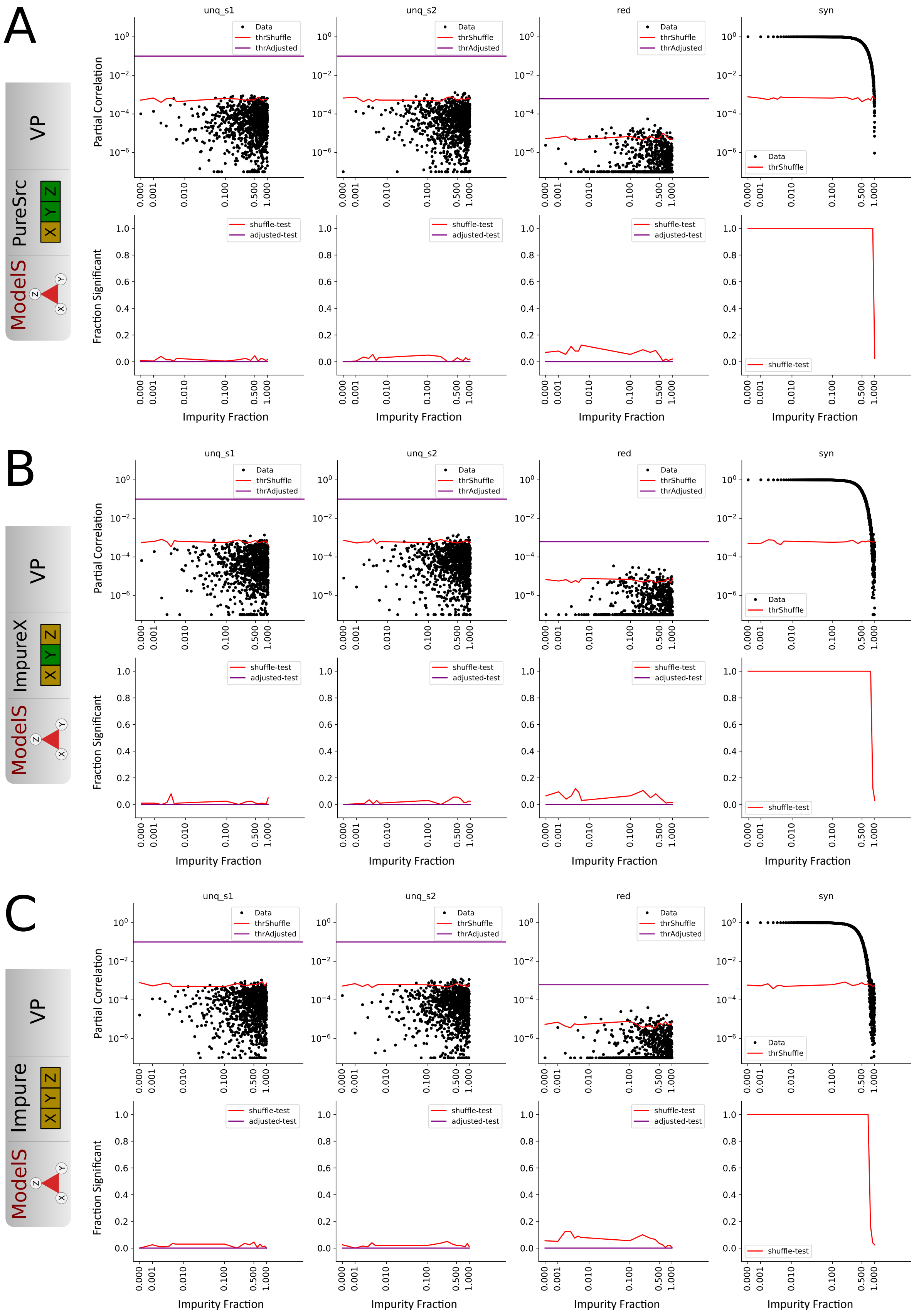


Figure 7: Variance Partitioning magnitude (top) and fraction of significant values (bottom) for ModelS and different observable models, as function of impurity fraction for $N_{tr} = 10000$. Red line denotes permutation testing threshold, and corresponding fraction of significant values. Purple line denotes the same for the adjusted conservative test. Columns in each figure denote information atoms $U(X \to Z|Y)$ and $U(Y \to Z|X)$, $R(X, Y \to Z)$ and $S(X, Y \to Z)$ respectively.

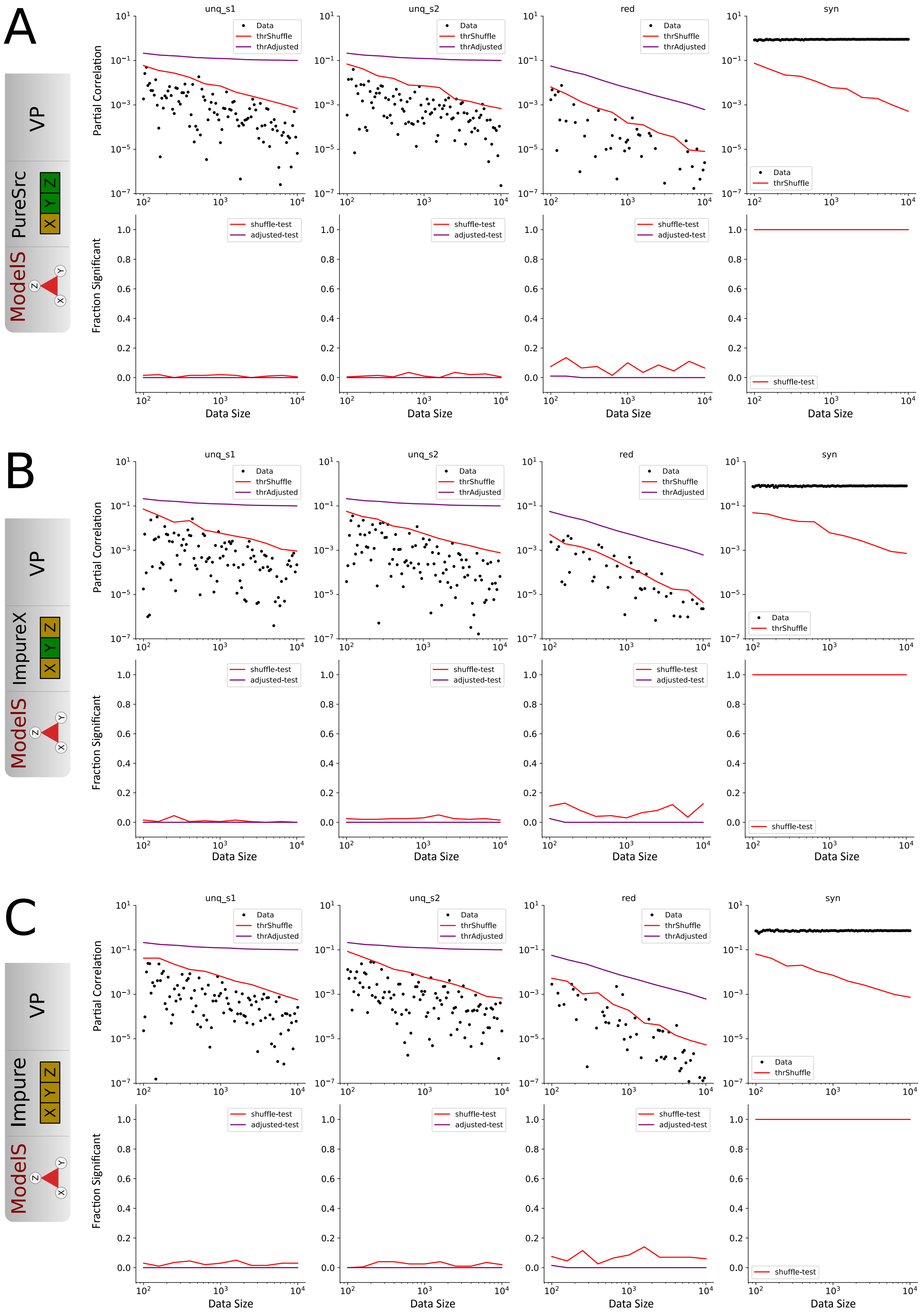


Figure 8: Variance Partitioning magnitude (top) and fraction of significant values (bottom) for ModelS and different observable models, as function of data size for fixed impurity fraction 0.25. Red line denotes permutation testing threshold, and corresponding fraction of significant values. Purple line denotes the same for the adjusted conservative test. Columns in each figure denote information atoms $U(X \to Z|Y)$ and $U(Y \to Z|X)$, $R(X, Y \to Z)$ and $S(X, Y \to Z)$ respectively.

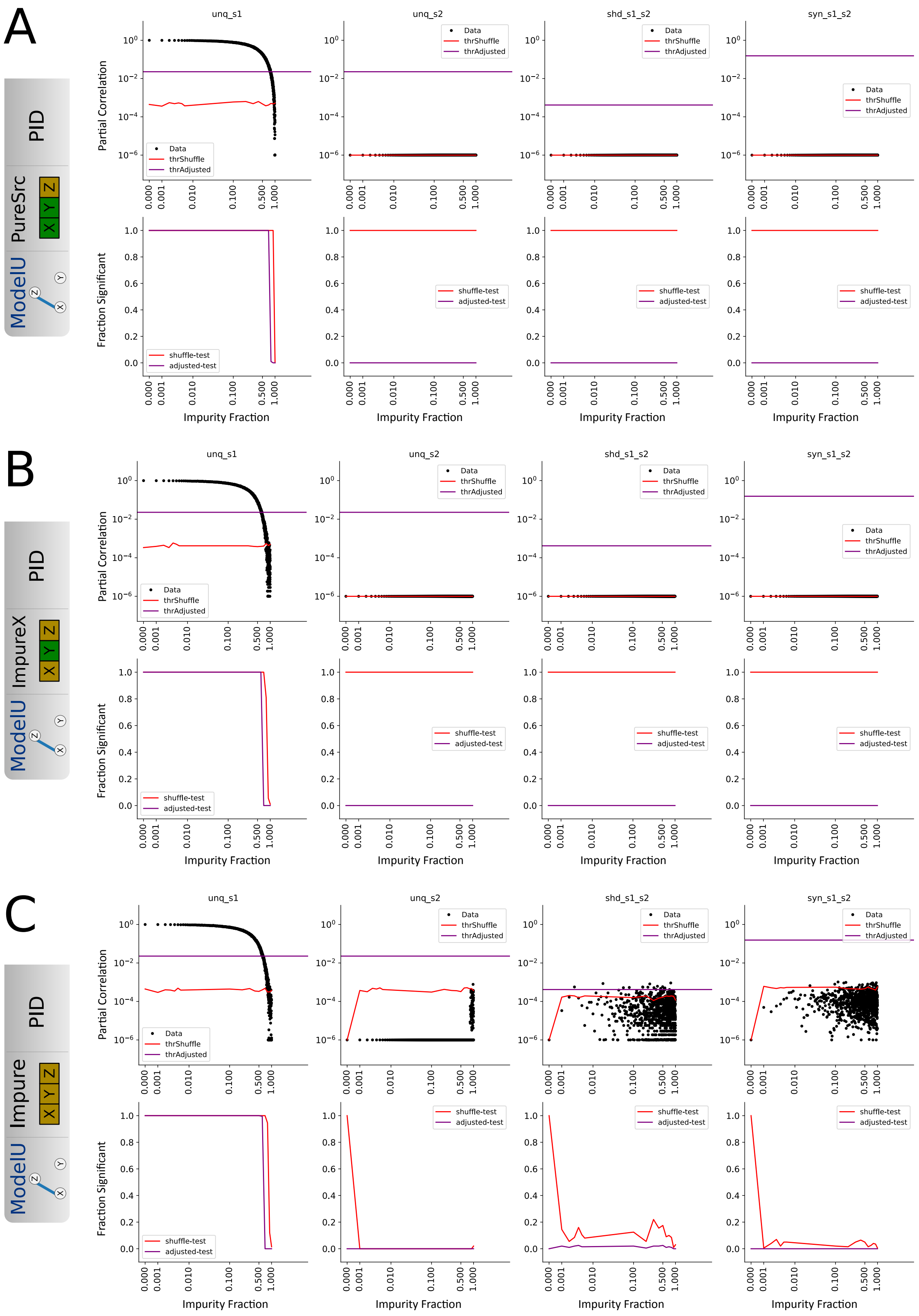


Figure 9: Partial Information Decomposition magnitude (top) and fraction of significant values (bottom) for ModelU and different observable models, as function of impurity fraction for $N_{tr} = 10000$. Red line denotes permutation testing threshold, and corresponding fraction of significant values. Purple line denotes the same for the adjusted conservative test. Columns in each figure denote information atoms $U(X \to Z|Y)$ and $U(Y \to Z|X)$, $R(X, Y \to Z)$ and $S(X, Y \to Z)$ respectively.

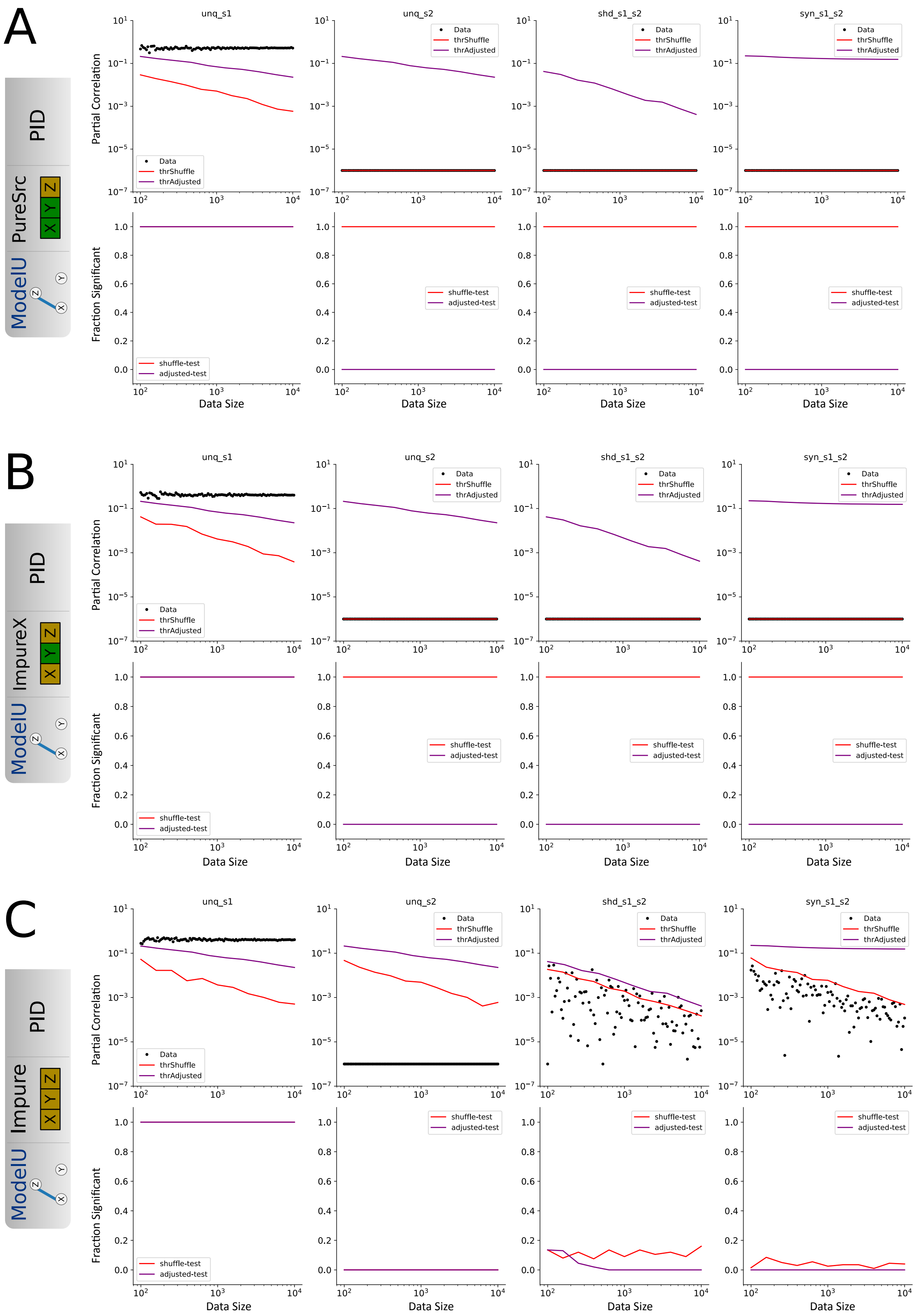


Figure 10: Partial Information Decomposition magnitude (top) and fraction of significant values (bottom) for ModelU and different observable models, as function of data size for fixed impurity fraction 0.25. Red line denotes permutation testing threshold, and corresponding fraction of significant values. Purple line denotes the same for the adjusted conservative test. Columns in each figure denote information atoms $U(X \to Z|Y)$ and $U(Y \to Z|X)$, $R(X, Y \to Z)$ and $S(X, Y \to Z)$ respectively.

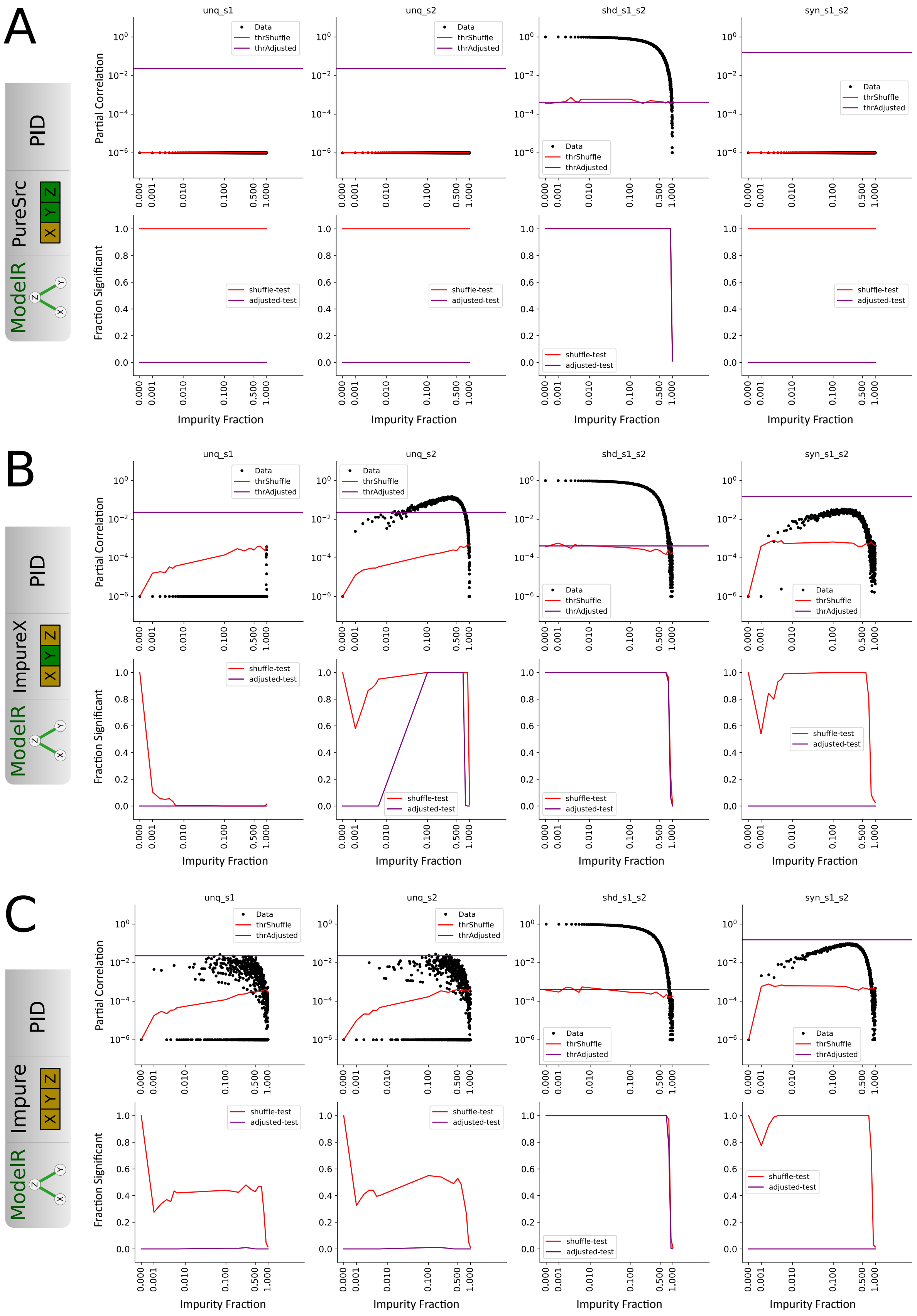


Figure 11: Partial Information Decomposition magnitude (top) and fraction of significant values (bottom) for ModelR and different observable models, as function of impurity fraction for $N_{tr} = 10000$. Red line denotes permutation testing threshold, and corresponding fraction of significant values. Purple line denotes the same for the adjusted conservative test. Columns in each figure denote information atoms $U(X \to Z|Y)$ and $U(Y \to Z|X)$, $R(X, Y \to Z)$ and $S(X, Y \to Z)$ respectively.

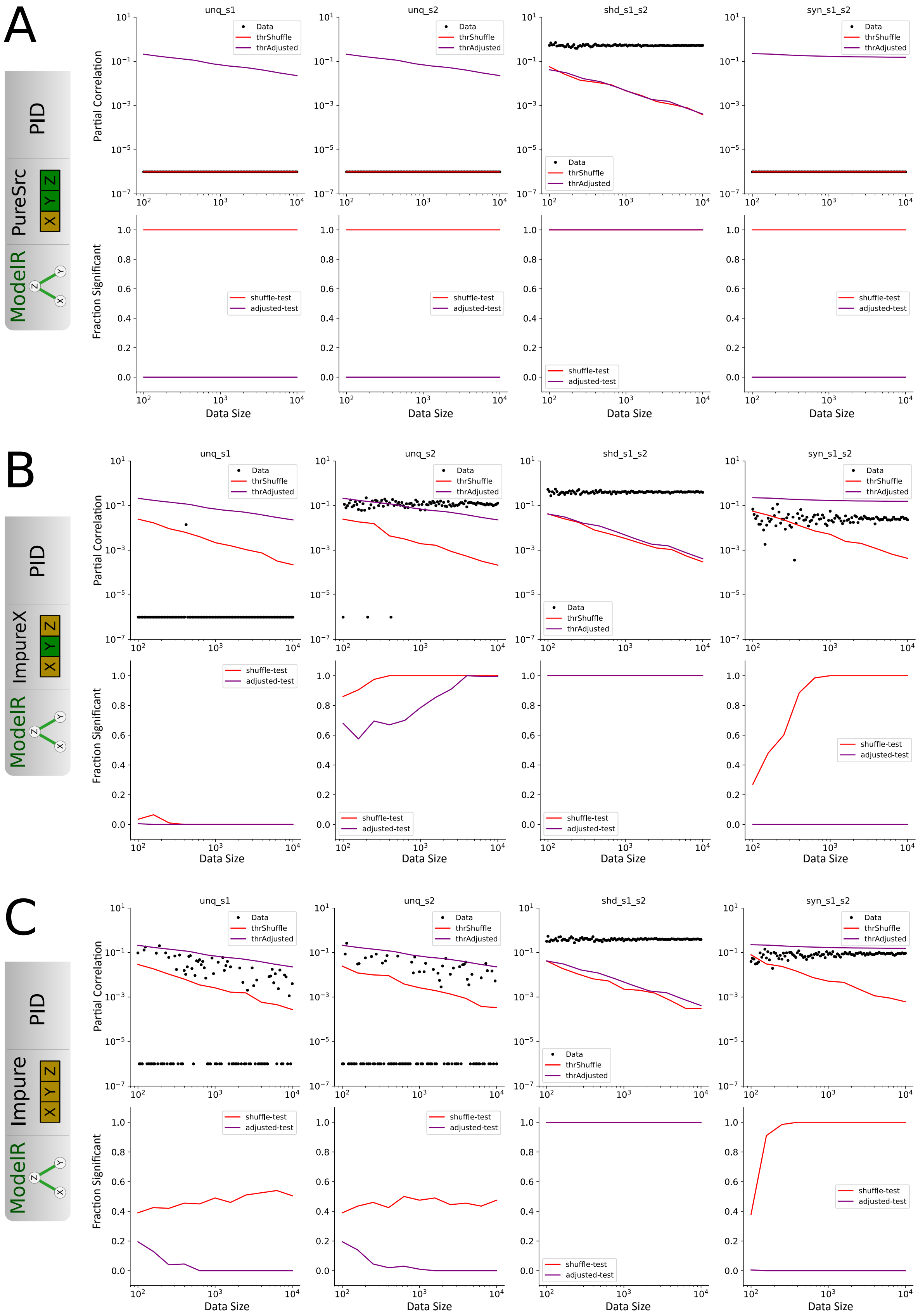


Figure 12: Partial Information Decomposition magnitude (top) and fraction of significant values (bottom) for ModelR and different observable models, as function of data size for fixed impurity fraction 0.25. Red line denotes permutation testing threshold, and corresponding fraction of significant values. Purple line denotes the same for the adjusted conservative test. Columns in each figure denote information atoms $U(X \rightarrow Z|Y)$ and $U(Y \rightarrow Z|X)$, $R(X, Y \rightarrow Z)$ and $S(X, Y \rightarrow Z)$ respectively.

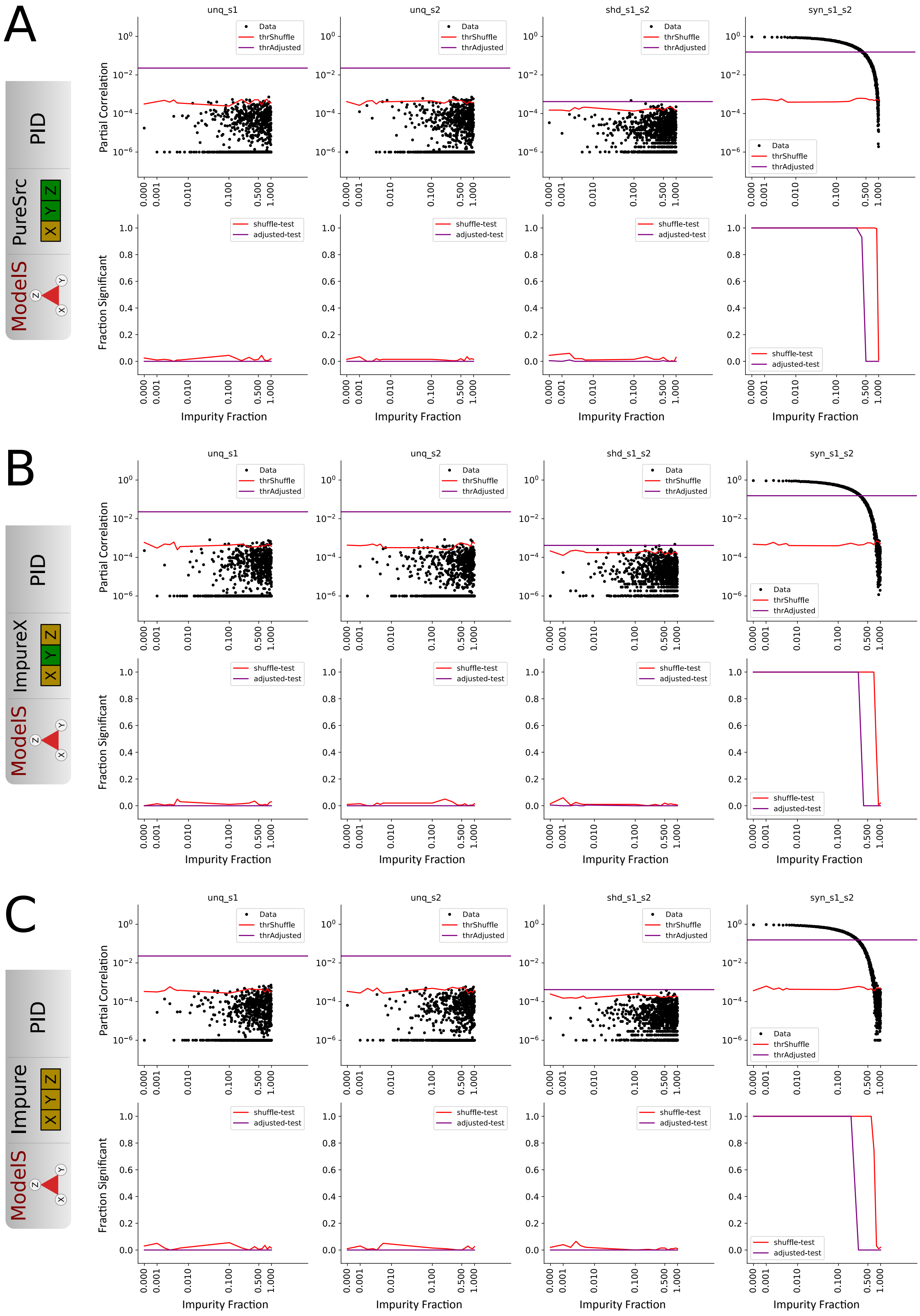


Figure 13: Partial Information Decomposition magnitude (top) and fraction of significant values (bottom) for ModelS and different observable models, as function of impurity fraction for $N_{tr} = 10000$. Red line denotes permutation testing threshold, and corresponding fraction of significant values. Purple line denotes the same for the adjusted conservative test. Columns in each figure denote information atoms $U(X \to Z|Y)$ and $U(Y \to Z|X)$, $R(X, Y \to Z)$ and $S(X, Y \to Z)$ respectively.

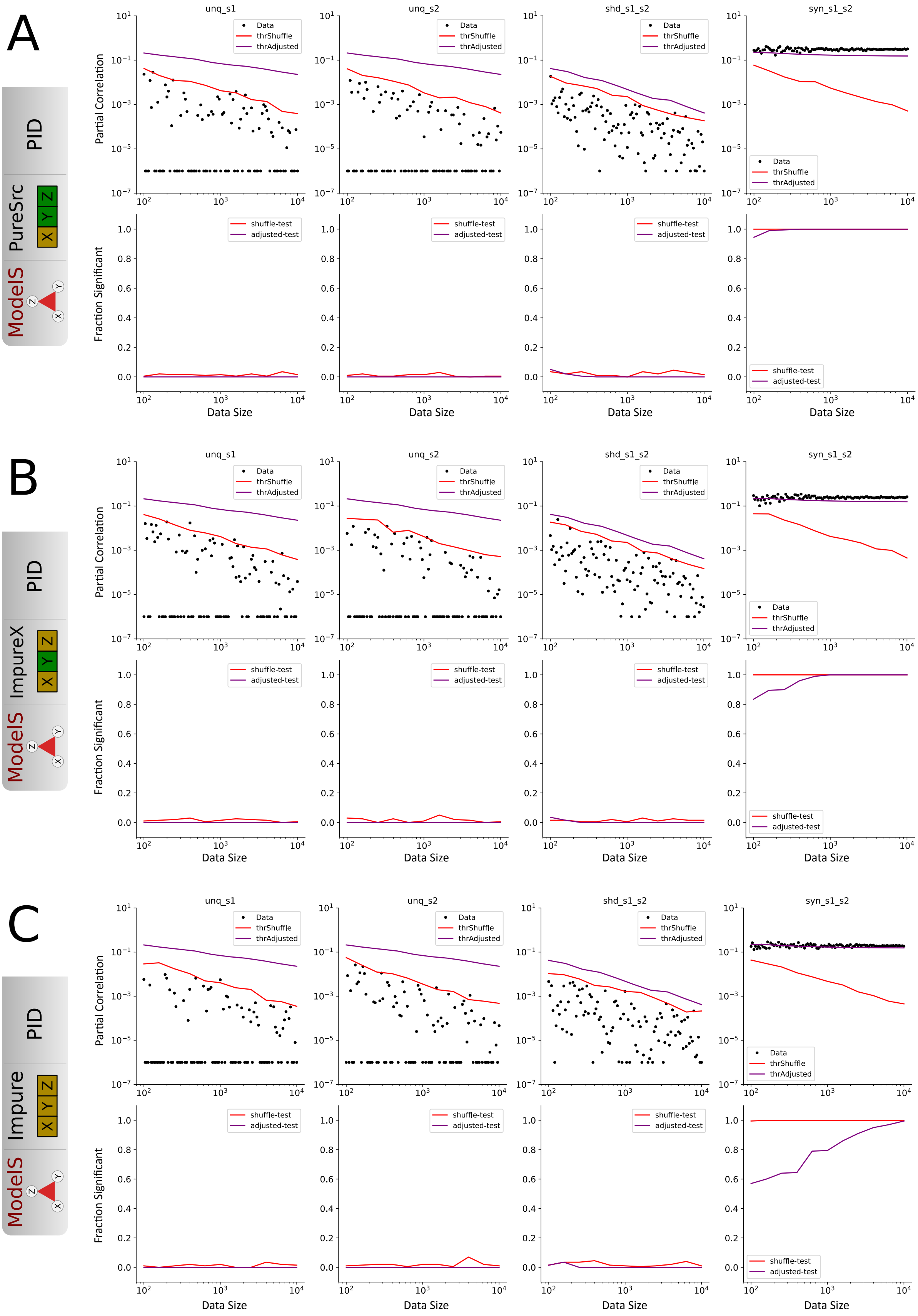


Figure 14: Partial Information Decomposition magnitude (top) and fraction of significant values (bottom) for ModelS and different observable models, as function of data size for fixed impurity fraction 0.25. Red line denotes permutation testing threshold, and corresponding fraction of significant values. Purple line denotes the same for the adjusted conservative test. Columns in each figure denote information atoms $U(X \to Z|Y)$ and $U(Y \to Z|X)$, $R(X, Y \to Z)$ and $S(X, Y \to Z)$ respectively.

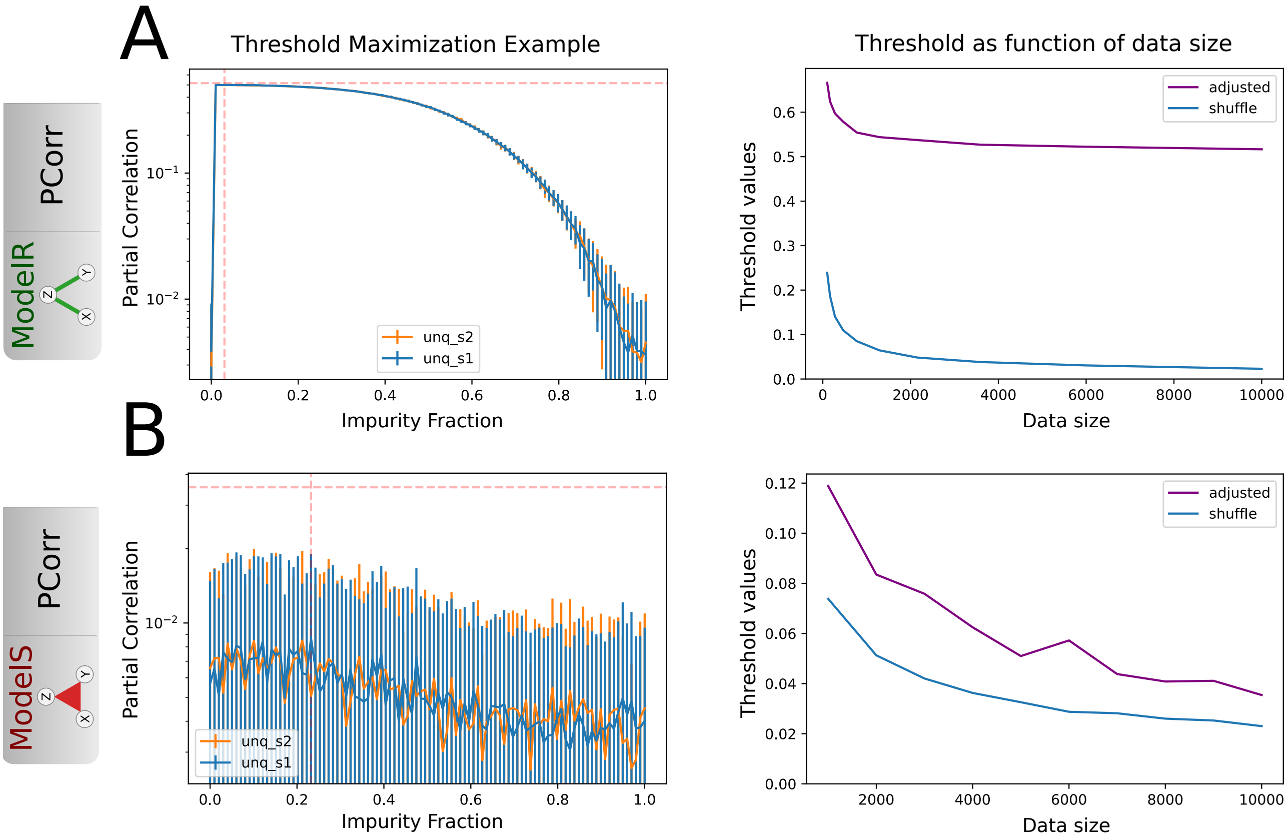


Figure 15: Adjusted testing threshold maximization for Partial Correlation A) for redundant adversarial model; B) for synergistic adversarial model. Left) False positive unique information atom distributions are plotted as function of impurity fraction for data size $N_{tr} = 10000$. Vertical dashed line denotes the impurity fraction corresponding to the highest threshold. Horizontal dashed line indicates the threshold (upper 1% quantile) at that impurity fraction. Right) Testing thresholds are plotted as function of data size. Blue denotes the threshold due to permutation-testing, purple denotes the adjusted testing threshold. For both models the adjusted threshold is above the permutation testing threshold. Thus, given either model as ground truth, permutation-testing would result in false positives.

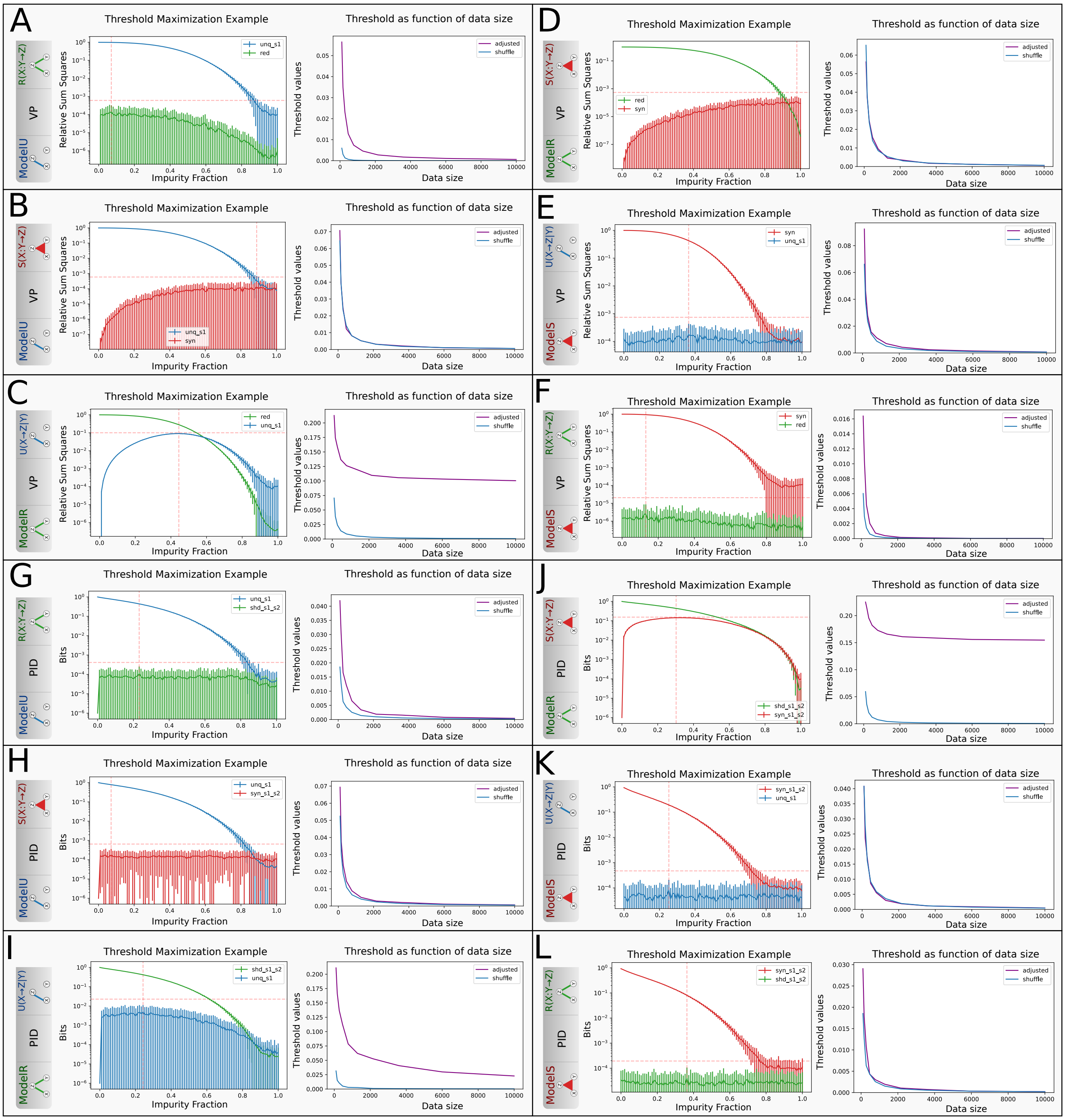


Figure 16: Adjusted testing threshold maximization for Variance Partitioning (A-F) and Partial Information Decomposition (G-L). Left and right plots same as in fig. 15. In the left figure, distribution of corresponding true positives is plotted alongside the false positives for comparison, but the false positives distribution is used to find the threshold. As described in the main text, in many combinations of ground truth model, observable model and the information atom the adjusted testing threshold does not exceed permutation-testing, suggesting there are no false positives for those combinations. This is mostly true for synergistic neuronal model, but is not true for redundant and unique models.

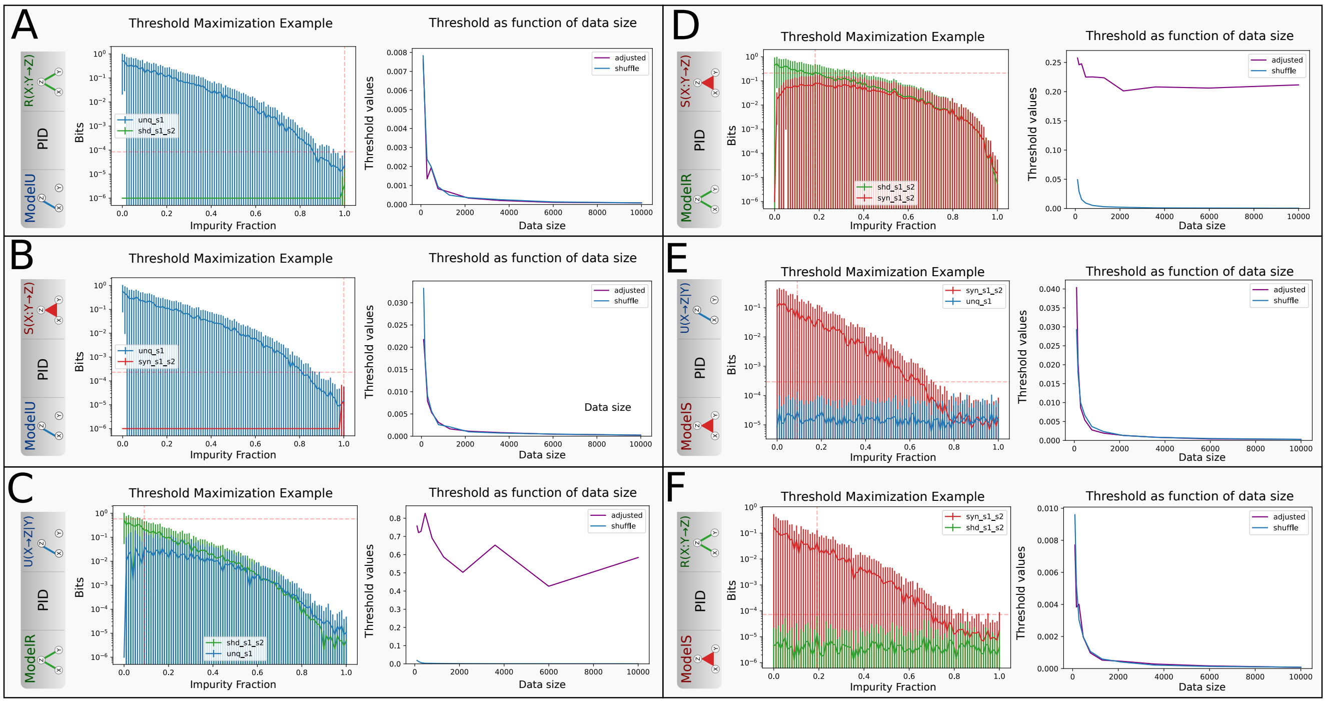

Figure 17: Adjusted testing threshold maximization for Partial Information Decomposition (PID) using discrete ground truth and observable models. Plot structure same as in fig. 16. PID exhibits false positives in unique and synergistic information atoms for redundant model, same as in the continuous case. However, unlike continuous case, it does not exhibit false positives in redundant atoms for unique model, suggesting that this may be purely an effect of binning. Nevertheless, the presence of false positives in discrete model excludes the hypothesis that they are all caused purely by the binning procedure.

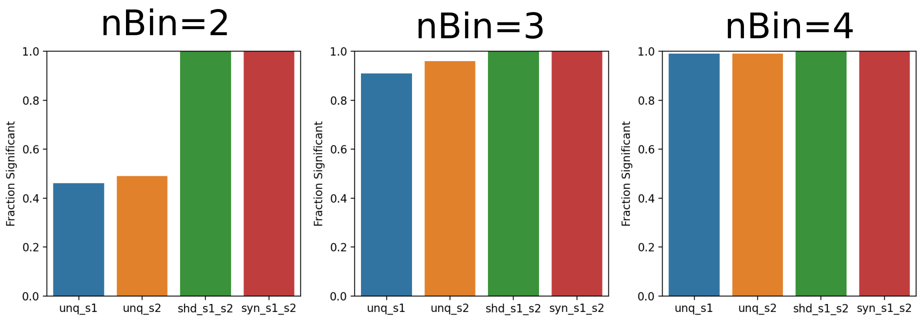


Figure 18: Effect of the data binning parameter. Plotted is the fraction of significant information atoms for the continuous Impure redundant model using Partial Information Decomposition, $N_{tr} = 10000$. Data for the three plots was binned using 2, 3 and 4 bins respectively. As can be seen, for all bin numbers, the number of true positive redundant atoms and false positive synergisitc atoms stays maximally at 100%. However, the number of false positive unique atoms gets progressively worse. This shows that the binning procedure alone can produce false positives in impure scenario.
